# A system-wide analysis of lipid transfer proteins delineates lipid mobility in human cells

**DOI:** 10.1101/2023.12.21.572821

**Authors:** Kevin Titeca, Antonella Chiapparino, Dénes Türei, Joanna Zukowska, Larissa van Ek, Mahmoud Moqadam, Sergio Triana, Inger Ødum Nielsen, Mads Møller Foged, Charlotte Gehin, Kenji Maeda, Theodore Alexandrov, Julio Saez-Rodriguez, Nathalie Reuter, Marco L. Hennrich, Anne-Claude Gavin

**Affiliations:** Department of Cell Physiology and Metabolism, University of Geneva, Geneva, Switzerland; European Molecular Biology Laboratory, EMBL, Heidelberg, Germany; Faculty of Medicine, and Heidelberg University Hospital, Institute for Computational Biomedicine, BioQuant, Heidelberg University, Heidelberg, Germany; Department of Chemistry, University of Bergen, Norway; Computational Biology Unit, Department of Informatics, University of Bergen, Bergen, Norway; Cell Death and Metabolism group, Center for Autophagy, Recycling and Disease, Danish Cancer Institute, Copenhagen, Denmark

**Author notes:** Current affiliation, Cellzome GmbH, a GlaxoSmithKline Company, Heidelberg, Germany (Marco L. Hennrich). Current affiliation: École Polytechnique Fédérale de Lausanne (EPFL), AI 1108 - Station 19, Switzerland. Equal contribution (Kevin Titeca, Antonella Chiapparino).

## Abstract

Lipid transfer proteins (LTPs) maintain the specialised lipid compositions of biological membranes, and many are associated with disease. In eukaryotes, they support organellar functions by transporting lipids between compartmentalised metabolic pathways. However, for the majority of the hundreds of human LTPs, the cargoes remain unknown. We combined biochemical, lipidomic and computational methods to characterize LTP-lipid complexes assembled *in cellulo* and in an *in vitro* biochemical assay. We identified bound lipids for about half of the LTPs analysed, and confirmed known cargoes, while discovering new ones for most LTP families. The data represents a systematic resource that captures the general principles of non-vesicular lipid transport in humans. The specificity of LTPs for lipids involves not only the recognition of specific head groups, but also of specific acyl chains. This selectivity defines lipid species within a lipid class with different metabolic or functional fates. The generalised ability of LTPs to form complexes with more than one class of lipids delineates new relationships between lipids and regulatory mechanisms that may contribute to the coordination of metabolism between different organelles. This work represents a resource and a framework for further analyses in different cell types, in pathological states or following various cellular perturbations.

---

Human cells generate thousands of different lipids^1^, the so-called lipidome, whose composition is adapted to cellular needs and functions. It directly influences biological processes, signalling efficiency and contributes to the establishment of cell identity and its functional specialization^2,3^. Importantly, all aspects of lipid function rely on their heterogeneous distribution in biological systems, where lipids accumulate locally and define the membranes of specific organelles or microdomains^4^. Maintaining optimal functional membrane composition involves compartmentalized lipid metabolism, associated with a variety of lipid sorting and transport systems, which can be provided, among other mechanisms, by lipid-transfer proteins (LTPs) ^5-9^. LTPs are structurally diverse, but many share a common mode of action: they bind and mobilise lipids into a hydrophobic pocket, forming stable, water-soluble protein-lipid complexes that isolate lipid cargoes from the aqueous phase and enable lipid transport within or between cells (reviewed in ^10-12^). In addition to their role in replenishing organelle lipid stores, some also act as chaperones presenting cargo to downstream protein effectors, such as enzymes, or as lipid sensors controlling cell signalling^8,10,13-15^.

LTP functions are essential and conserved in all kingdoms of life (reviewed in ^16^). There are at least 131 LTPs in humans, whose dysfunction is often associated with various diseases^12,17,18^. However, we only understand the function of a small number of them and for the majority the identity of their cargoes remains elusive. As they have never been systematically studied, we still lack a complete mapping of LTP-mediated lipid transport pathways; how they overlap, cooperate and act synergistically. In previous proof-of-principle studies in a model eukaryote, *Saccharomyces cerevisiae*, we demonstrated the feasibility of systematic analyses of lipid transfer^6^. We therefore set out to characterize human LTP-lipid complexes assembled under physiological conditions in a human cell line and in an *in vitro* biochemical assay. Due to its unprecedented scale, the resulting resource significantly expands our understanding of non-vesicular lipid transport in humans.

## A systematic resource for the study of human non-vesicular lipid transport

The lipid binding properties of LTPs have not yet been systematically studied and the general principles of their specificity and functions have remained elusive. Here, we measured the ability of human LTPs to mobilize specific lipid cargoes, i.e. to load them and carry them in aqueous environments. To this end, we have adapted affinity purification methods coupled with lipidomics to characterize soluble LTP-lipid complexes^6^, and applied them to two complementary approaches (Supplementary methods). We measured the ability of affinity-tagged LTPs overexpressed in HEK293 cells to associate stably with lipids in the physiological context of a human cell (Fig. 1a; *in cellulo*). We also studied the ability of recombinant LTPs, expressed in *Escherichia coli*, to extract lipids from artificial membranes composed of lipids extracted from bovine liver and porcine brain (Fig. 1a; *in vitro*).

In total, 101 human LTPs were cloned, and we were able to express 86 in HEK293 cells and 71 in *E. coli* (Fig. 1a). We successfully purified 110 LTP-lipid complexes assembled *in cellulo* or *in vitro* (counting the redundancy of two screens) by affinity and size exclusion chromatography (SEC) (Fig. 1a). The SEC fractions were analysed by sodium dodecyl sulfate-polyacrylamide gel electrophoresis (SDS-PAGE) and liquid chromatography-tandem mass spectrometry (LC-MS/MS)- or high-performance thin-layer chromatography (HPTLC)-based lipidomics (Supplementary methods; Supplementary Data S1A, S1B and S1C; Extended Data Table 1) ^6^. To filter out non-specific background, we matched LTP abundance in SEC fractions with lipid abundance. Only lipids identified in all fractions containing LTPs and showing an elution profile (i.e., changes in abundance in fractions) similar to that of LTPs were considered as potential lipid binders (Supplementary methods). The work consisted of more than 600 LC-MS/MS runs, and the analysis of the resulting large datasets required the development of semiautomatic pipelines, incorporating quality filtering and manual processing of the spectra (Fig. 1b and Extended Data Fig. 1) (Supplementary methods). To ensure the accuracy and depth of identification of the different lipid species (for example to characterize the composition of their fatty acid chains), we measured the experimental fragmentation and elution patterns of series of lipid standards to define fragmentation rules (Extended Data Tables 2A and 2B), which we combined with spectral information from the databases METLIN, SwissLipids and LIPID MAPS^19-21^. We also considered evaluating the overall quality of the LC-MS/MS-based analyses^22-25^. As expected from a reverse-phase LC system, lipids with long fatty acids (i.e., more hydrophobic) had longer retention times than those with short fatty acids, whereas lipids with unsaturated fatty acids eluted more rapidly than their saturated counterparts (Extended Data Fig. 2a). The fact that we often identified lipid species in the form of different adducts and in different MS ionization modes also increased confidence in their identification (Extended Data Fig. 2b).

We identified lipid cargoes for 45 of the LTPs that we could affinity purify (Fig. 1a Extended Data Table 1). The datasets cover nine of the ten LTP families, with up to 10 representatives for the lipocalins^10^. While the majority of LTPs analysed were successfully expressed and purified in both expression systems (HEK293 and *E. coli*), only six extracted lipids in both assays (Fig. 1a). This shows that the two approaches are complementary. The *in cellulo* screen is more likely to capture lipid cargoes of LTPs whose assemblies require a cellular machinery. However, it is also limited to one biology, that of exponentially dividing HEK293 cells, where specific pathways and functions are inactive (see examples below). The *in vitro* approach offers a broader scope, as LTPs are exposed to a wide variety of lipids, but we will miss the lipid cargoes of LTPs whose assemblies require a cellular machinery. The integration of both *in cellulo* and *in vitro* methods provide systematic data on LTP-lipid complexes and represents an important resource.

## General properties of the LTP-associated lipidome

This study represents a comprehensive analysis of the lipid-binding properties of human LTPs, and should help define the general principles of lipid handling by these proteins. We first investigated whether the lipids recognised by the analysed LTPs differ from the total lipidome, i.e., whether the LTP system shows preferences for certain lipid attributes or properties. To do this, we measured the total lipidome of HEK293 cells cultured under the same conditions by LC-MS/MS-based lipidomics and compared it with the lipidome mobilised by LTPs (Fig. 2 and Extended Data Table 5).

We observed well-known patterns, for example the enrichment of lipids in fatty acids with an even number of carbons, in particular 32, 34, and 36 (counting the two acyl chains), both in the total cellular lipidomes and those associated with LTPs (Fig. 2a,b). Interestingly, the presence of odd-numbered species is somewhat more frequent in LTP-lipid complexes assembled *in vitro* (Fig. 2a and Extended Data Fig. 2c). This likely reflects their exposure to a wider range of lipids, including those of the expression host, *E. coli*, where odd numbered lipids are more frequent^26^. In line with this notion, these exclusively assembled *in vitro* complexes often contain lipids that are abundant in bacteria but not in humans (e.g. phosphatidylglycerol) (Fig. 2b and Extended Data Fig. 2c), suggesting that some may have been assembled in *E. coli*^6^. These *in vitro* data suggest that they insert biochemically into the binding pocket, and are therefore informative as to the nature and extent of this pocket’s binding.

We also observed some interesting general trends. The LTPs studied in both screens preferentially mobilised glycerophospholipids with shorter fatty acids, whereas we did not observe a similar trend for sphingolipids, which seem to have more complex selectivity patterns (Fig. 2b) (see below). The shorter fatty acids might facilitate extraction from membranes because of their reduced lateral hydrophobic interactions. Similarly, LTPs also showed a preference for glycerophospholipids and sphingolipids bearing only one to two sites of unsaturation in their fatty acids (i.e., carrying one or two double bounds on the acyl group and long-chain base) (Fig 2c). These lipid species can cause deep membrane defects, a phenomenon that can contribute to the membrane lipid uptake^27^. In contrast, polyunsaturated fatty acids (PUFAs), and fully saturated fatty acids cause only superficial or no defects, respectively, and may be more difficult to extract from membranes^27^.

Importantly, the LTP associated lipidome sometimes reveals lipid species that were barely, if at all, detectable in the total lipidome of HEK293 cells, such as rare, long-chain ceramide species with 46 or 48 carbons (acyl group plus long-chain base) (Fig. 2b and Extended Data Table 5) (see below). The enrichment for rare and low-abundant lipids, unlikely to come from nonspecific contamination of the total lipidome, provides an additional level of confidence in the dataset. More interestingly, it also suggests that within the major lipid classes, aliphatic chain lengths and saturation define lipid pools that are differentially accessible to the LTP system and that may experience different metabolic fates (see below).

## An LTP-lipid interactome map reveals new routes of lipid transfer

The screens confirmed many of the previously known LTP cargoes (Fig. 3a,b), knowledge that can be used to assess the quality and the complementarity of the two screens (Extended Data Table 1). For example, some of the known associations between LTPs and lipids, such as phosphatidylcholine with STARD2 and STARD10, are observed in both screens, while others are screen-specific. This highlights the unique ability of each approach to detect a different set of complexes. Among the known LTP-lipid complexes observed only *in cellulo* are those whose assembly requires a cellular machinery. For example, the retinol-binding proteins, RBP1 and RBP4 formed complexes with vitamin A only in the cellular context. This is likely because their assembly requires active metabolism and transmembrane transport systems, which are absent in the *in vitro* assay^28^.

In other cases, LTPs load their known cargoes only in the *in vitro* biochemical assay, i.e., where they escape spatial or temporal cellular controls and possible contextual imbalances in cellular lipid availabilities^29^. Members of two evolutionary distinct LTP families that transport phosphatidylinositols, the precursors of phosphatidylinositol phosphates (PIPs) synthesis, illustrate this situation. Some members of the CRAL-TRIO family, including SEC14L2, are known to transport phosphatidylinositols from the ER for phosphorylation into specific PIPs that accumulate in the membranes of various organelles, contributing to organelle identity and maintenance^30-32^. The PITPNA, PITPNB and PITPNC1 belong to the phosphatidylinositol transfer protein (PITP) family and participate in the so-called phosphatidylinositol cycle. They transfer phosphatidylinositol from the ER to the plasma membrane to replenish the phosphatidylinositol pool after signalling and phospholipases C-mediated hydrolysis of PI(4,5)P_2_^33^. Although phosphatidylinositol is abundant in HEK293 cells, *in cellulo* we did not find it associated with PITPs, but with SEC14L2 (Fig. 3a,c and Extended Data Fig. 3). This is probably due to the fact that SEC14L2-phosphatidylinositol complexes contribute to important functions in dividing cells that need to duplicate their organelles (such as the HEK293 cells used here), whereas phospholipase signalling is probably not active in this cellular context, so PITP-phosphatidylinositol complexes do not assemble. PITPs formed complexes with phosphatidylinositol only in the *in vitro* assay. It is interesting that PITPs primarily form complexes with PI(38:4), an arachidonic acid-containing species (Fig. 3c), which is known to be predominant in the phosphatidylinositol cycle^34^. This demonstrates that PITPNA, PITPNB and PITPNC1, even in the absence of a cellular machinery, prefer arachidonic acid-containing phosphatidylinositol species, and that this ability is likely to contribute to the metabolic routing of lipids with specific fatty acids to the phosphatidylinositol cycle^34^.

We also identified lipids for nine LTPs with no previously known ligands: SEC14L6, ATCAY, BNIPL, BPIFB2, SEC14L5, TTPAL, LCN15, SCP2D1, and HSDL2 (Fig. 3a and Extended Data Table 1). For example, *in cellulo* BPIFB2 forms complexes with bismonoacylglycerolphosphate (BMP), a rare lysosomal lipid whose biosynthesis pathway has only recently been characterized, suggesting a possible role in these processes^35,36^. Hydroxysteroid dehydrogenase like 2 (HSDL2) is a peroxisomal multi-domain LTP^37^ (it has a peroxisomal targeting signal) that bound to triacylglycerol *in vitro* (Fig. 3a and Extended Data Table 1), supporting the idea that it plays a role in peroxisomal triacylglycerol transport and metabolism. Consistent with this hypothesis, the knockdown of HSDL2 leads to the accumulation of lipid droplets^38,39^ and several of the known HSDL2 interactors are peroxisomal and involved in β-oxidation (Extended Data Fig. 5a,b). OSBPL9 belongs to the family of oxysterol binding proteins. Members of this family are known to exchange phosphatidylinositol-4-phosphate with sterols or with phosphatidylserine between cellular membranes^6,40-42^. OSPBL9 binds to phosphatidylserine *in cellulo* (Fig. 3a and Extended Data Table 1), consistent with the prediction of a phosphatidylserine-binding motif in its lid region^6^.

We even identified lipids, such as BMP or triacylglycerol (see above), that are novel in the LTP transport system. Another example is diacylglycerol, which *in vivo* formed cellular complexes with SEC14L2, a lipid chaperone acting as a lipid-presenting scaffold for several lipid kinases, e.g. the phosphatidylinositol 4-kinase^43^, the phosphatidylinositol 3-kinase or an as yet unidentified alpha-tocopherol (vitamin E) kinase^44^. As expected, *in cellulo* SECL14L2 formed complexes with phosphatidylinositol (see above and Extended Data Fig. 3), but also with a vitamin E-like lipid that is not based on a squalene backbone nor contains fatty acids. Interestingly, SEC14L2 also bound rare diacylglycerol species (DAG(32:1) and DAG(34:2)) *in cellulo* (Extended Data Fig. 3). This association with diacylglycerol may provide a molecular mechanism for SEC14L2-dependent coordination of phosphatidylcholine biosynthesis (via CDP-choline, i.e., using diacylglycerol as a precursor) with SEC14L2-mediated stimulation of phosphatidylinositol 4-kinase activity^45^. In this perspective, it is intriguing that diacylglycerol is also the substrate of a family of ten diacylglycerol kinases, which produce phosphatidic acid. Overall, these examples illustrate that integrating *in cellulo* and *in vitro* approaches to study LTP-lipid complexes in a systematic manner uncovers many new potential lipid cargoes, which should inspire and trigger future work.

## Lipid transfer proteins can bind to more than one class of lipids

A few LTPs, such as members of the OSBP, PITP or the CRAL-TRIO families, are known to accommodate more than one class of lipids and this property contributes to the coordination of metabolism between different organelles. Some directly exchange lipids, for example, cholesterol in the ER for PI4P in the Golgi^8,13,40^. For others, the binding of a sensor or regulatory lipid in a deep pocket initiates various chaperonin functions and the presentation of another lipid (i.e. phosphatidylinositol) to downstream enzymes (e.g., kinases) ^45^. Here, we demonstrate that many (22) of the (39) LTPs studied are capable of forming complexes with more than one class of lipids, suggesting that these regulatory mechanisms represent general attributes of LTP functions (Fig. 4a,b).

We reasoned that lipid pairs mobilised by the same LTPs should share physiological relationships. Therefore, we evaluated the links of these pairs in other lipidomics data, specifically whether they are more frequently co-regulated and co-localised than any other random lipid pairs. For this, we integrated data describing changes in lipid abundance following systematic knockdown of enzymes from sphingolipid metabolism^2^ (Fig. 5a), and scored all lipid pairs for their co-regulation. In this scoring system, a “+1.0” denotes perfect co-regulation (lipids always change abundance in the same way), whereas a “-1.0” denotes mutual exclusion (lipids always change oppositely). We also exploited a large spatial metabolite database, METASPACE, which consists of thousands of tissue sections analysed by imaging mass spectrometry (Fig. 5b). Each pixel in these images corresponds to one mass spectrometry analysis (Supplementary methods) ^46^. Finally, we also included a dataset capturing the lipidome of affinity-purified organelles, to refine lipid co-localisation to subcellular levels (unpublished data) (Fig. 5c). For all lipid pairs in these datasets, we determined their co-occurrence by the Manders’ overlap coefficient^47^ (Supplementary Methods)(Fig. 5b,c). In this system, “+1.0” and “0.0” represent lipid pairs that always or never co-localize, respectively. Interestingly, we observed that the lipid pairs mobilised by the same LTPs are significantly more co-regulated upon metabolic perturbation (Fig. 5a) and co-localised (Fig. 5b,c) than would be expected from random sets of lipid pairs (Extended Data Table 4). These trends remain significant, when only lipid classes are considered, i.e. when excluding closely related lipid species or lipid sub-classes from this analysis (Fig. 5). Overall, this supports the idea that LTPs can be organised into functional cellular networks linking distinct organelles and contributing to the coordination of metabolism between different cellular compartments.

## Interaction map reveals new principles of sphingolipid transport

We identified the expected cargoes for LTPs known to mediate the intracellular movement of several sphingolipids^5,48,49^ (Extended Data Table 1). CERT binds ceramides (Fig. 3a), consistent with its contribution to their transport from the ER^5,50,51^, where they are made, to the Golgi, where they are metabolised to sphingomyelin, ceramide-1-phosphate or glycosphingolipids^52-54^. GLTPD1 forms complexes with ceramide 1-phosphate^55^, and GLTP with glycosphingolipids^54,56-58^ (hexosylceramides) (Figs. 3a), which is known to trigger their transport to the plasma membrane. In addition to these well-known examples, we have seen that a lipocalin, LCN1, binds to sphingomyelin (Fig. 3a), a lipid previously not known to be part of this LTP system, as well as to phosphatidylcholine, another choline-containing lipid (Fig. 3a). LCN1 is the major lipid-binding protein in tears, a fluid indeed highly enriched in phosphatidylcholine and sphingomyelin^59^.

The acyl chain length of ceramides is tightly controlled, and there are six ceramide synthases in mammals that produce ceramides with specific acyl chain distributions^48^. We noted that *in cellulo* CERT exhibits interesting patterns of specificity for distinct ceramide species, which appears to define different pools with distinct metabolic fates (Fig. 6a). For example, it forms complexes with rare, saturated and very long dihydroceramides and phytoceramides - 46 and 48 carbons (long chain base and fatty acid length)(Fig. 6a). Dihydroceramides are metabolic intermediates, produced directly by ceramide synthases, that can be converted to ceramides or phytoceramides by the addition of a double bond or a hydroxyl group at position C-4, respectively^60-62^. This suggests a novel role for CERT in the early sorting of very long-chain dihydroceramides, i.e. prior to conversion to ceramides, for the production of phytoceramides, which are believed to play important structural function in the maintenance and stability of ordered membrane microdomains^63^.

Interestingly*, in cellulo*, the 40- to 42-carbon ceramides produced by ceramide synthases 2 and 3^64^, although abundant in HEK293 cells, are not mobilised by CERT (Fig. 6a), which is consistent with previous *in vitro* data^51^. These 40- to 42-carbons structures are particularly abundant in the pool of glycosylated ceramides, i.e., those that have acquired hexoses and that are handled by GLTP (Fig. 6a), consistent with the idea that they were transported to the Golgi by a mechanism independent of CERT^5,65^. In contrast, CERT efficiently mobilised the 32- to 36-carbons ceramides. These structures are abundant in the sphingomyelin and ceramide 1-phosphate pools that are handled by GLTPD1 and GLTP, respectively (Fig 6a). To further investigate the role of CERT in ceramide sorting, we measured the impact of its gain of function on cellular lipidomes (Fig. 6b). We observed that over expression of CERT in HEK293 resulted in an increase in cellular sphingomyelin and ceramide 1-phosphate, and a decrease in hexosylceramides. Overall, these data support the idea that CERT is part of a ceramide triage system, thus defining three pools of ceramides with different metabolic fates.

We also observed novel lipid binders for the known sphingolipid transporters, suggesting the existence of novel regulatory mechanisms. For example, GLTPD1, consistently formed complexes not only with its cognate cargo, ceramide 1-phosphate^55^, but also with sphingomyelins both *in cellulo* and *in vitro* (Figs. 3a and 6a). Sphingomyelins did not appear to be actively transported by GLTPD1 in *in vitro* transfer assays^55^ (data not shown). The function of these complexes with sphingomyelins remains elusive. They may play regulatory roles, for example, by coordinating ceramide 1-phosphate metabolism (phosphorylation/dephosphorylation) to local sphingomyelin requirement and availability. The other example is CERT, which, in addition to ceramide species also associates with phosphatidylcholine (Fig. 3a). Binding of phosphatidylcholine (palmitoyl-oleoyl-phosphatidylcholine, POPC) to CERT through insertion of one fatty acid tail into its lipid binding pocket was also observed in molecular dynamics simulations of the CERT-START domain with membranes using an all-atoms model^66^. Simulations of CERT-phosphatidylcholine adducts in an aqueous environment in the presence and absence of the cargo ceramide resulted in both cases in the formation of stable complexes. In these simulations the hydrophobic cavity accommodates the two lipid tails from the phosphatidylcholine, also in the presence of ceramide (Fig. 6c,d). Phosphatidylcholine is one of the two substrates (along with ceramide) of sphingomyelin synthase, which catalyses the transfer of the phosphocholine head of phosphatidylcholine to ceramide to produce sphingomyelin. This is consistent with the idea that CERT could act as a chaperone, a notion already proposed for the CRAL-TRIO family^43^, bringing both substrates at the same location, and coupling or coordinating their transfer to their transformation into sphingomyelin.

## Discussion

By combining two complementary approaches, this study systematically highlights lipid specificities for the numerous human LTPs. This combination elucidates both lipid binding as a function of cellular context and identifies lipids that accommodate into lipid-binding pockets. This is an extensive resource that should enhance our understanding of the pathways linking different parts of lipid metabolism to signalling. The extend of this research has identified general and specific features for the lipids observed with LTPs, including previously unknown unifying patterns. Comparison of the whole-cell lipidome with that associated with LTPs revealed that not all lipid species are easily manipulated by LTP systems. These showed a general preference for short, single or double unsaturated lipids, known to create deep defects in membrane^27^, which should contribute to their extraction from membranes. Different lipid species within the same class will therefore have different mobility in cellular systems. The work has provided novel concepts and important resources, as well as new bioinformatics pipelines for matching protein and lipid elution profiles, and for identifying protein-bound lipids, with the establishment and use of highly detailed spectral rule sets. They are relevant to the wider field of lipid-protein interaction research, and provide a starting point for extending such analyses to different cell types or following different cell perturbations or stimulations.

**Figure 1:**
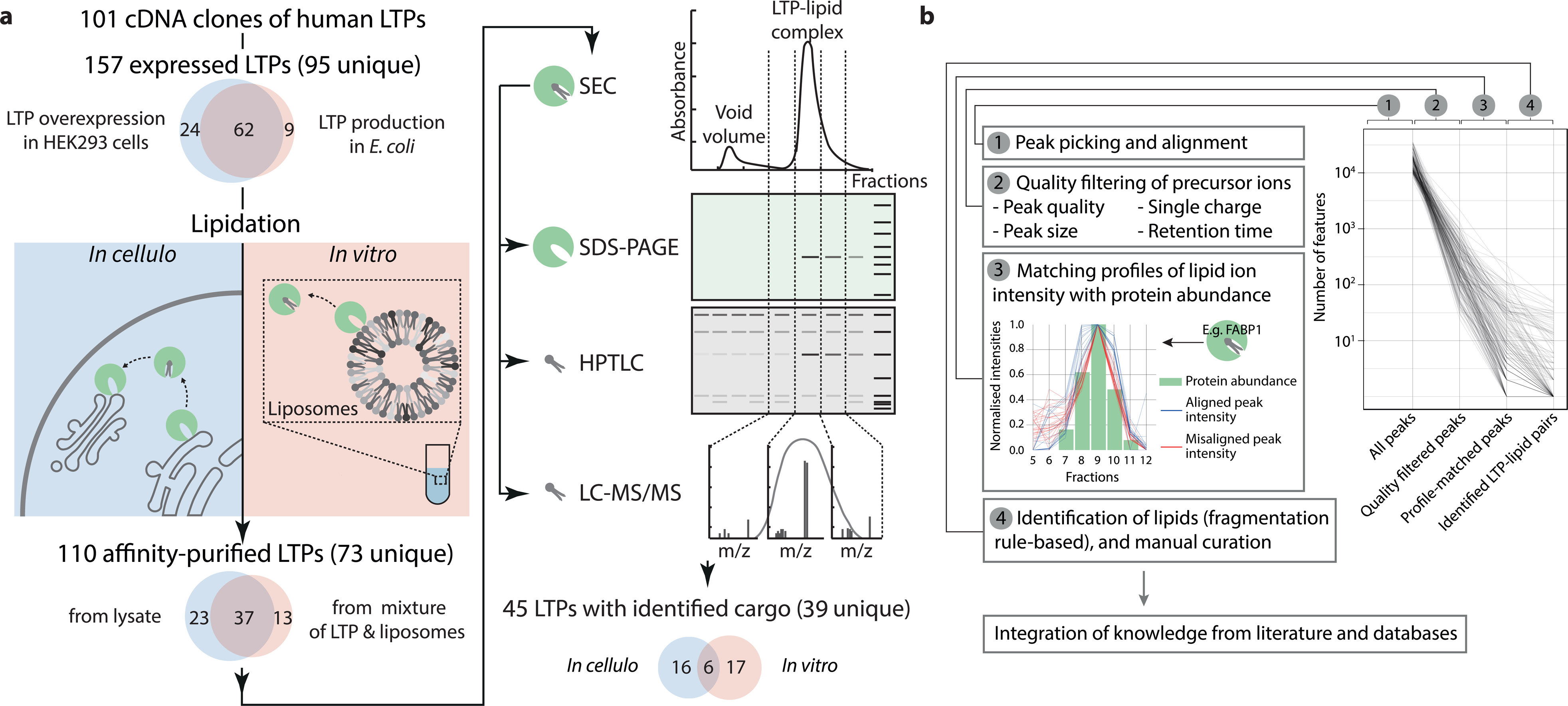
Synopsis of the screens. (a) Overview of the analysis steps of the *in cellulo* and *in vitro* screens and the associated success rates. (b) Data analysis pipeline overview.

**Figure 2:**
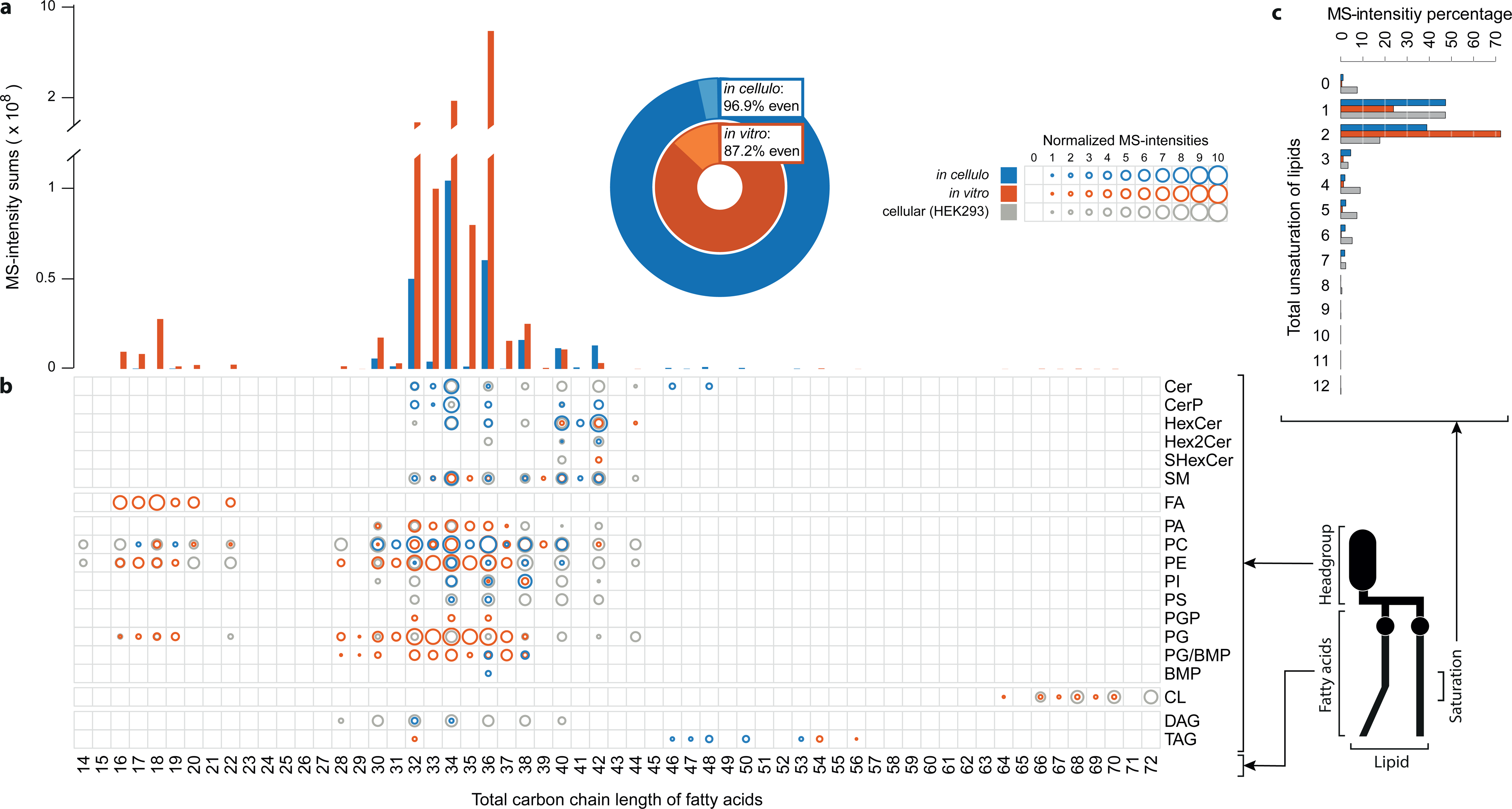
Characterization of the mobilised lipidome of the screens, with a focus on different lipid parts and on the lipids observed with mass spectrometry. (a) Total fraction of the intensities with an even versus odd total carbon chain length of the fatty acyls for the *in cellulo* and *in vitro* screens, and distribution of the total sum of the intensities for all total carbon chain lengths of the fatty acyls in the *in cellulo* and *in vitro* screens. (b) Distribution of normalised intensities of all observed combinations of headgroups and total carbon chain lengths of the fatty acyls. Comparison of the *in cellulo* (blue) and *in vitro* (orange) distributions of the mobilised lipids with the lipids present in the whole HEK293 cells (grey) (Extended Data Table 5) (vitamins were not included). (c) Distribution of total summed intensities for each of the different degrees of fatty acyl unsaturation of lipids identified in the *in cellulo* screen, *in vitro* screen and HEK293 cells. Expressed as a percentage of total summed intensities for each lipidome.

**Figure 3:**
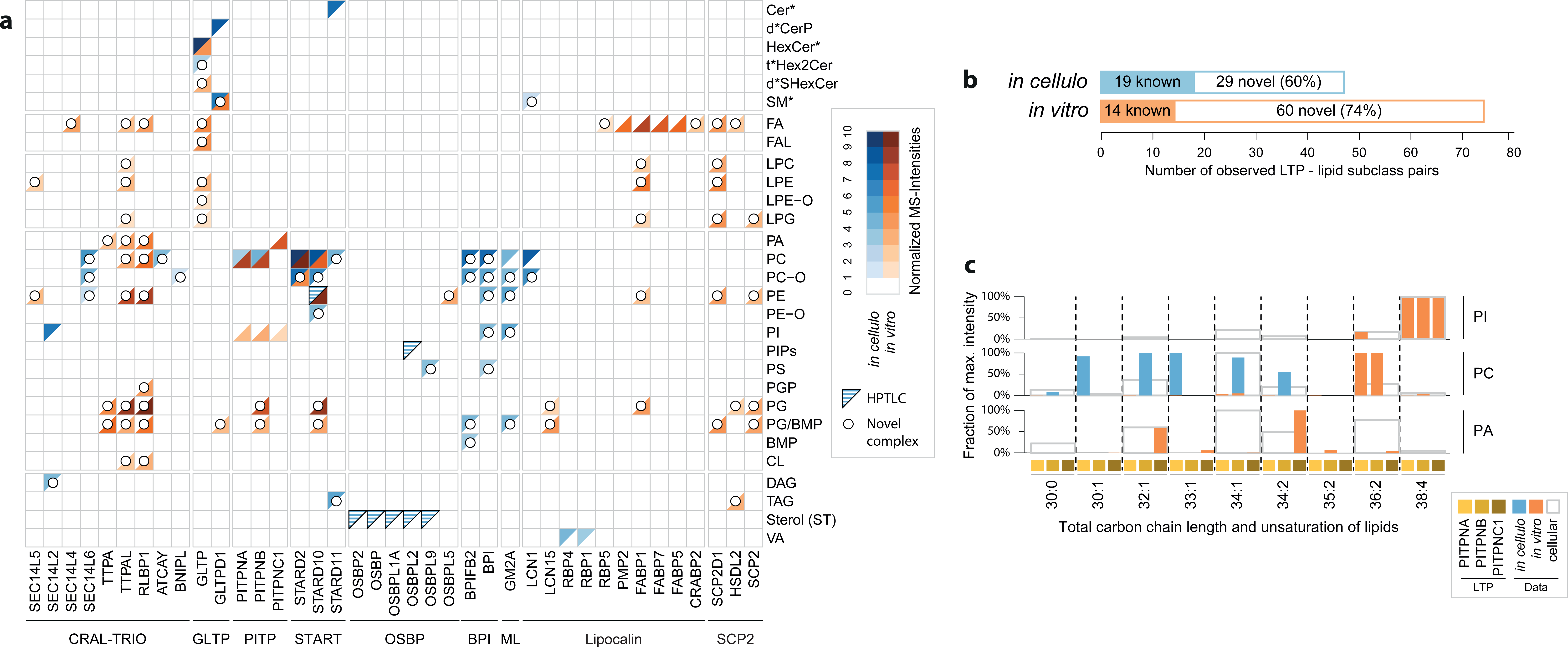
Overview of the lipids mobilised by the LTPs. (a) Normalised intensities for all observed LTP-lipid subclass combinations for the *in cellulo* and *in vitro* screens, and an overview of their individual novelties. HPTLC-based observations are only represented when no mass spectrometry data was available (black bordered blue triangles). The LTPs were seriated by the presence and location of all of their protein domains, and grouped along the LTP family lines (Extended Data Fig. 4). (b) General number of known and novel LTP – lipid subclass pairs for the *in cellulo* and *in vitro* screens. (c) Distribution of lipid species for PI, PC and PA for the PITP-family of LTPs. The observed intensities of the individual species are normalised to the maximum observed for the specific LTP – lipid subclass pair. The *in cellulo* (filled blue) and *in vitro* (filled orange) distributions for the LTPs are compared with distribution of the lipids present in the whole HEK293 cells (Extended Data Table 5) (grey border).

**Figure 4:**
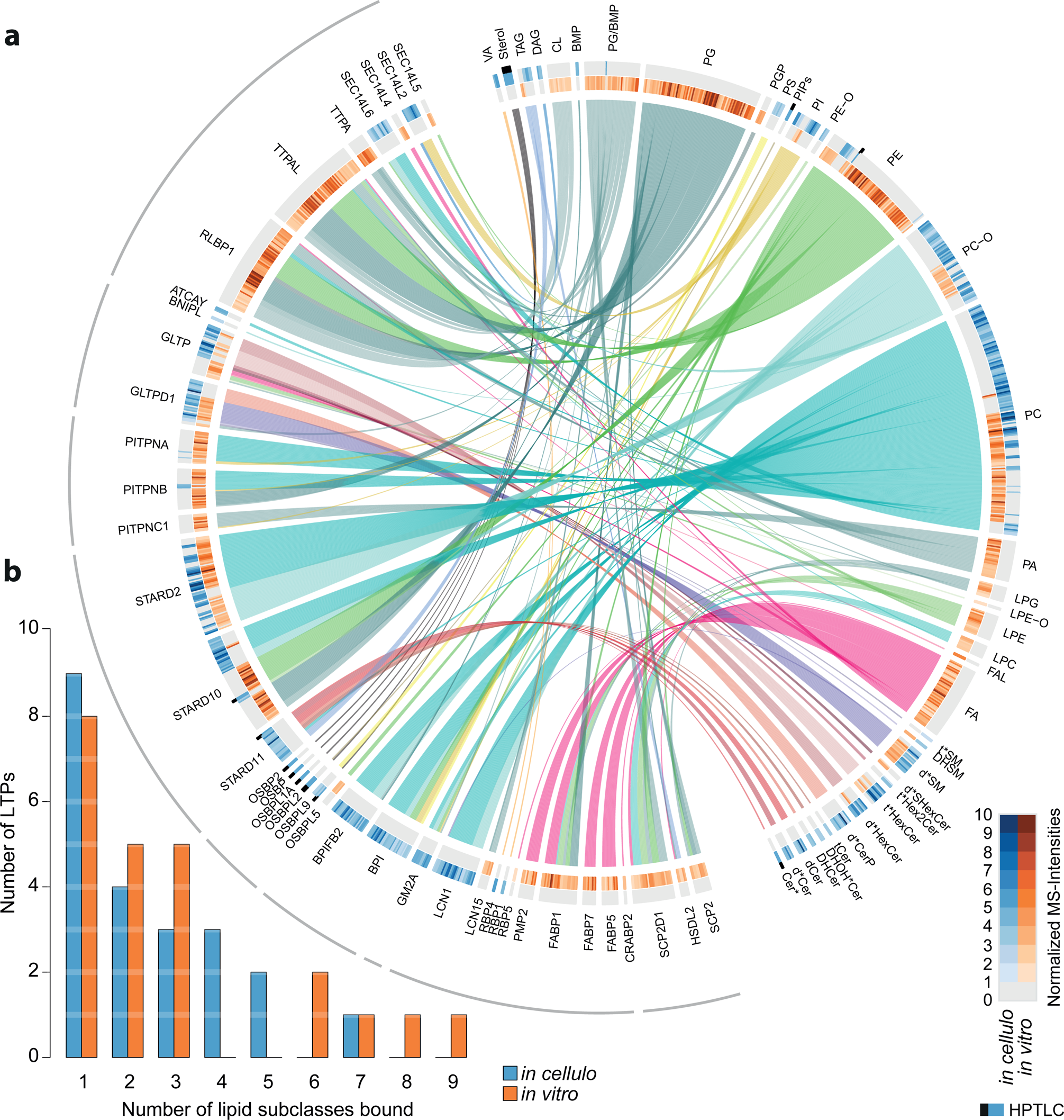
Most LTPs mobilize multiple lipid subclasses, and most lipid sub-classes are also mobilised by multiple LTPs. (a) Overview of all LTP – lipid sub-class pairs for the *in cellulo* and *in vitro* screens. HPTLC-based observations are only represented when no mass spectrometry data was available (black squares on top of blue squares). The LTPs were seriated by the presence and location of all of their protein domains, and grouped along the LTP family lines (Extended Data Fig. 4). (b) General distribution of the number of LTPs binding different numbers of lipid subclasses for the *in cellulo* and *in vitro* screens.

**Figure 5:**
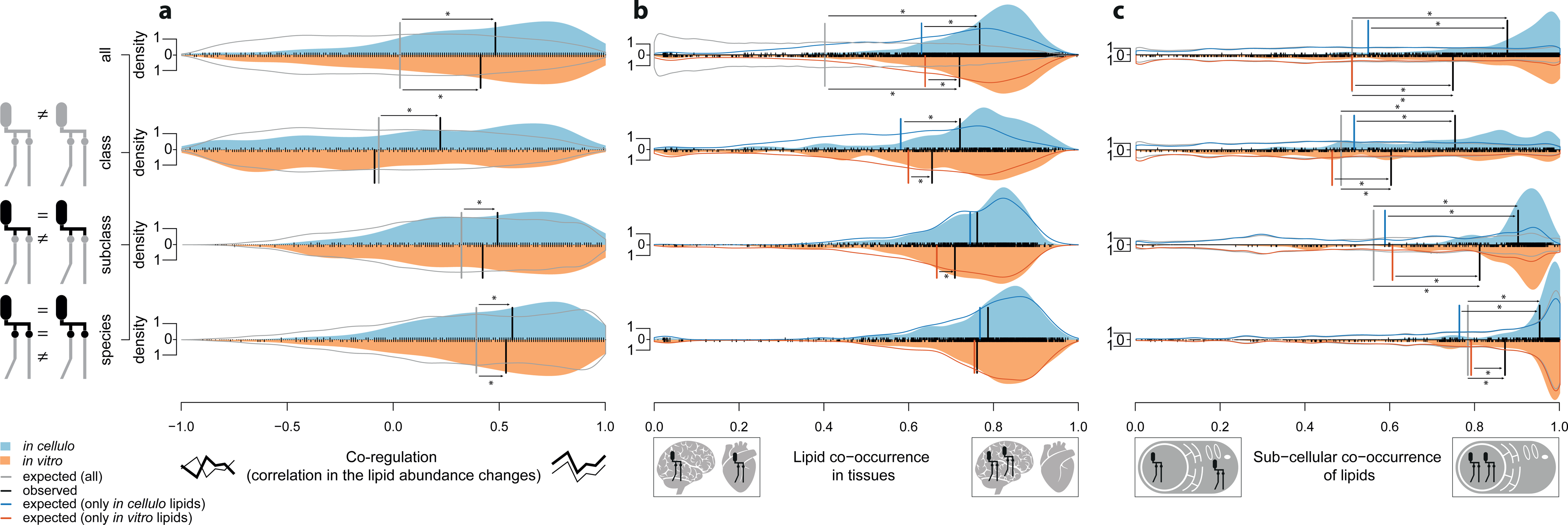
Co-mobilised lipids by LTPs are also co-regulated and co-localized. Vertical lines in the graphs represent medians. Significant shifts of the observed medians are indicated by a star (see Extended Data Table 4 for significance calculations). The expected distribution lines in grey are always based on all entries in the reference database, while the expected distribution lines in colour are only based on the observed lipid classes from the reference database by either the *in cellulo* (blue) or *in vitro* (orange) screens. The densities with Gaussian kernels use bandwidth selection according to^67^. An overall density of 1 would correspond to a uniform distribution. All densities have the same surface area for all comparisons within each panel. (a) Comparison of the co-mobilization of lipids by the LTPs with the co-regulation of these lipids, according to the correlation of the lipid abundance changes upon cellular perturbations as described in^2^. (b) Comparison of the co-mobilization of lipids by the LTPs with the Manders’ co-occurrence of these lipids in tissue section in the METASPACE database^46,47^. (c) Comparison of the co-mobilization of lipids by the LTPs with the Manders’ co-occurrence of these lipids in sub-cellular fractions^47^.

**Figure 6:**
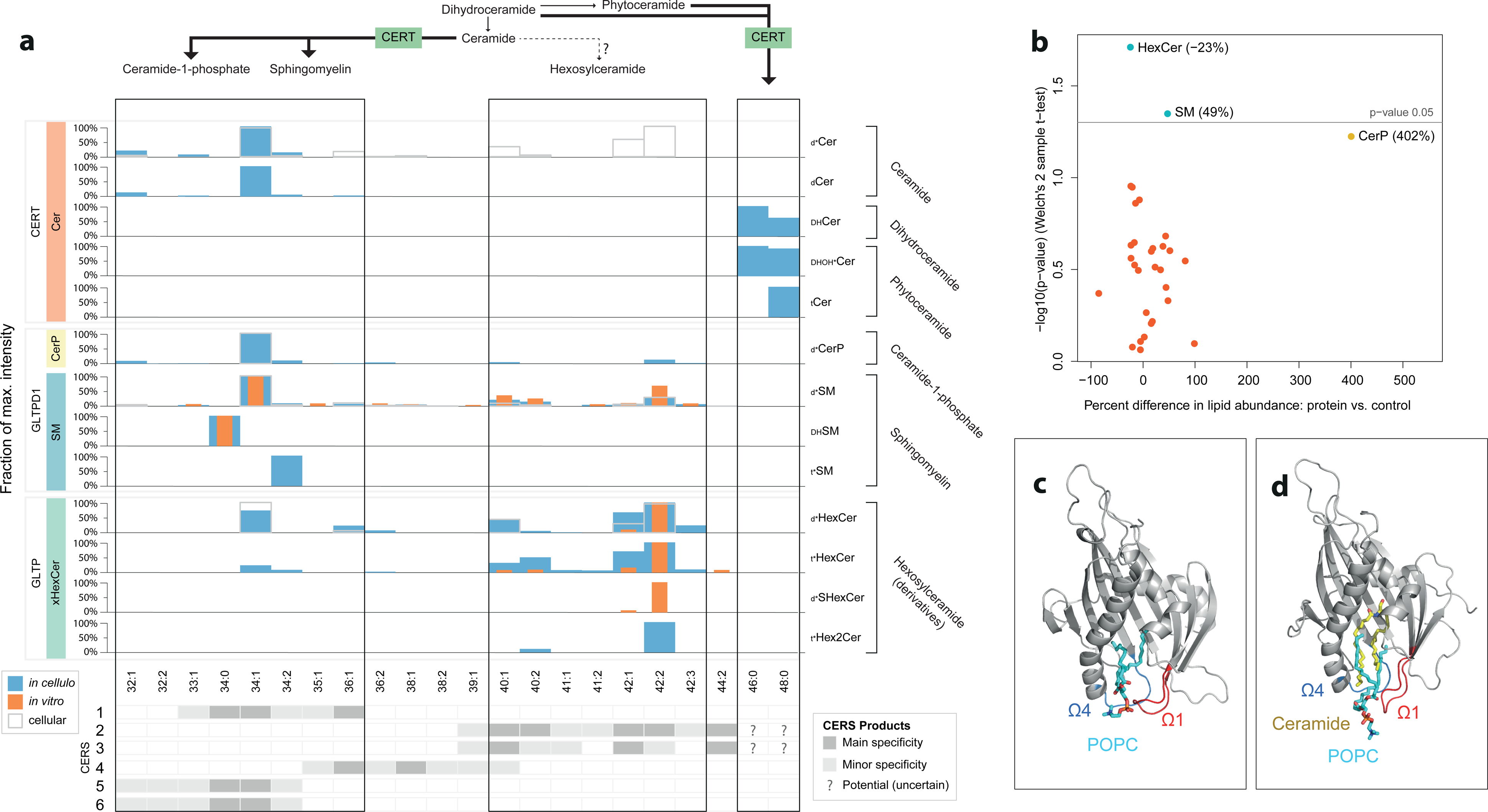
Comparison of the species distributions of the sphingolipid mobilizing LTPs for the *in cellulo* (full blue) and *in vitro* (full orange) distributions of the mobilised lipids with the lipids present in the whole HEK293 cells (Extended Data Table 5) (grey border). The observed intensities of the individual species are normalised to the maximum observed for the specific LTP – lipid subclass pair. The data of all hexosyl-containing sphingolipids were integrated into xHexCer. The bottom highlights the specificity of each ceramide synthase (CERS) ^64^. (a) Overview of the sphingolipid total acyl chain lengths transported by CERT, GLTPD1 and GLTP. At the top: mobilization by CERT of short-to-medium length d ceramides and long saturated non-d ceramides. Ceramide mobilization by CERT is absent for acyl chain lengths downstream of CERS2,3 and potentially 4. In the middle: broad SM mobilization by GLTPD1 in a similar pattern to the mobilised ceramide-1-phosphate (Cer1P). At the bottom: mobilization by GLTP of hexosyl-containing species with a preference for longer total carbon chain lengths. (b) The effect of overexpression of CERT in HEK293 cells on the cellular lipid class abundances. (c) and (d) Molecular dynamics simulations show that the START domain can accommodate a phosphatidylcholine (palmitoyl-oleoyl-phosphatidylcholine, POPC) in its hydrophobic cavity, both in the absence (c) or the presence (d) of ceramide (Supplementary Methods). Ceramide coloured yellow, POPC coloured cyan, Ω1 coloured red and Ω4 coloured blue. START domain is shown as cartoon and coloured grey.

The study identified many new lipid-binding partners, opening up new avenues of research. The role of lipid headgroups in the specificity and mechanism of lipid transfer has long been recognised. We have observed, for example, that LCN1 binds lipids with a choline head group, while the type of fatty acid linkage (ether or ester), their length or even the class of lipid (sphingomyelin or phosphatidylcholine) appear to play little role in the recognition mechanisms. Due to the unbiased nature of our assays, where LTPs select lipids from whole lipidomes comprising all species (which has rarely been done before), we can start assessing specificity beyond lipid head groups. These data suggest that within a lipid class, LTPs have preferences for specific species, and that this selectivity defines lipid pools with different metabolic fate and/or cellular functions.

Most LTPs associate with multiple lipid classes, while this was previously only known for a few of them, this work now supports the notion that this is a general attribute shared by many LTPs. Integration with different types of lipidomics datasets further validated the functional relevance of co-transported lipid pairs. Interestingly, this is significant not only for related lipid species (i.e. lipids with structural similarity or belonging to the same metabolic pathways), but also for lipid classes with no structural and metabolic relatedness. This confirms the idea that lipid co-transport represents an important, but hitherto underestimated, mechanism for the organization and integration of lipid metabolic pathways and the maintenance of organellar membrane function.

## Acknowledgements

We are grateful to the EMBL Metabolomics core facility; the Proteomics platform of the University of Geneva, and the READS platform of the University of Geneva, for expert helpWe thank Lipotype for its expertise in lipidomic analysis and Vladimir Rybin, EMBL, for his help with protein expression and biophysical characterization. The thank Howard Riezman, University of Geneva, and Emmanuel Varesio, University of Geneva, for inspiring discussions and for sharing their lipidomics expertise. We are grateful to Bernd Klaus, EMBL, and Matt Rogon, EMBL, for sharing their expertise in bioinformatics and statistical data analysis and to Vladimir Rybin, EMBL, for expert help with protein purification. We are grateful to Enric Mila Vilalta who ran the *in vitro* lipid binding assay, and Laura Alvarez Aguilar and Michal Varga for generation of in house generated LTPs cDNA libraries and subsequent cloning in mammalian vectors. We thank Martin Beck and his group, in particular Amparo Andres-Pons, for sharing expression vectors and know-how on human cell culture and inducible systems. We also thank members and former members of ACG’s groups at the Department of Cell Physiology and Metabolism, University of Geneva and European Molecular Biology Laboratory, EMBL for continuous discussions and support, especially, Emeline Galster (University of Geneva). ACG acknowledge the financial support of the Louis-Jeantet foundation. KT was supported by the EU Marie Skłodowska-Curie Actions project 843407, LipTransProMet.

## Author contribution

ACG, AC, MLH designed the research; KT, DT, ACG, TA and JSR designed the data integration and analysis strategy; AC, JZ, LvE and CG conducted the molecular biology, cell biology and biochemistry experiments; MLH established the lipidomics platform with the help of AC; MLH, AC, IØN, KM conducted the mass spectrometry analyses and with KT interpreted their data; KT, DT and ST performed the bioinformatic analysis; MM and NR performed and interpreted the molecular dynamics experiment; KT and ACG wrote and all authors reviewed the manuscript.

