## Supplementary material for "A system-wide analysis of lipid transfer proteins delineates lipid mobility in human cells": Supplemetary Figures

Supplementary Figure 1: Broader overview of the steps described in the synopsis of figure panel 1b.

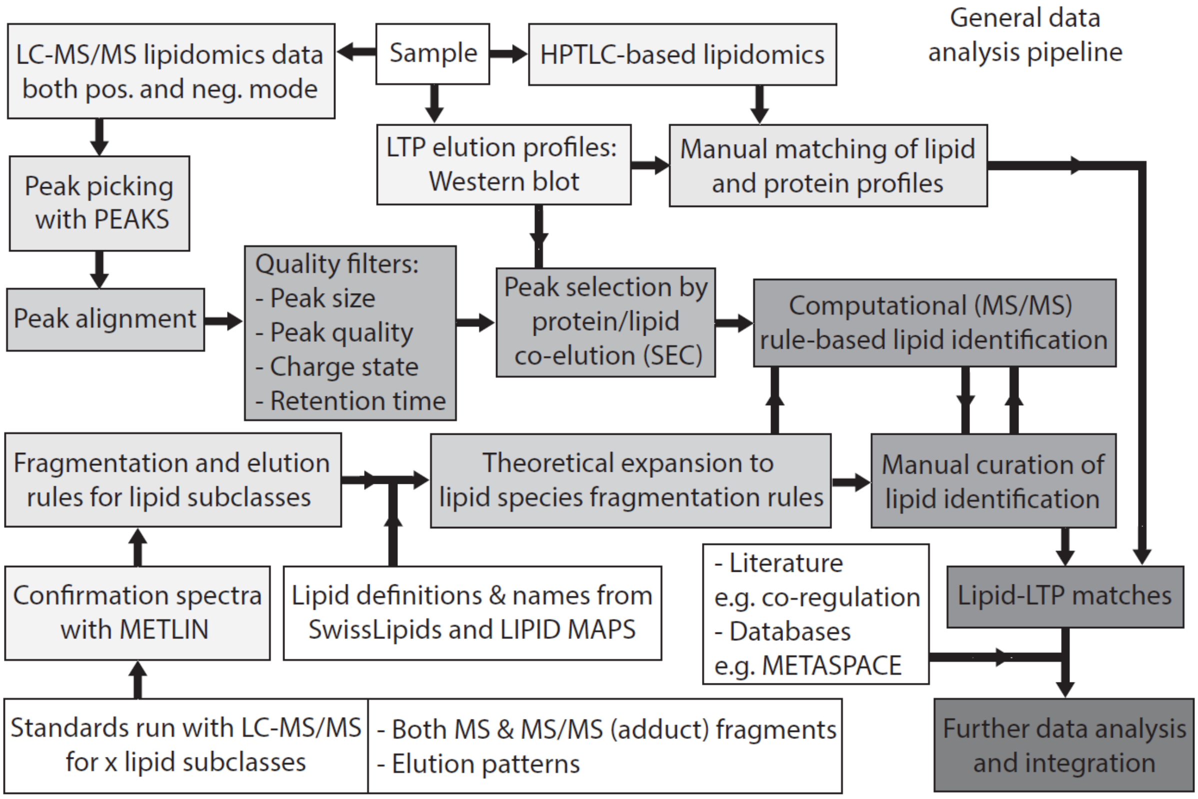


Supplementary Figures S2A: Quality control by comparison of lipid carbon chains vs. retention times

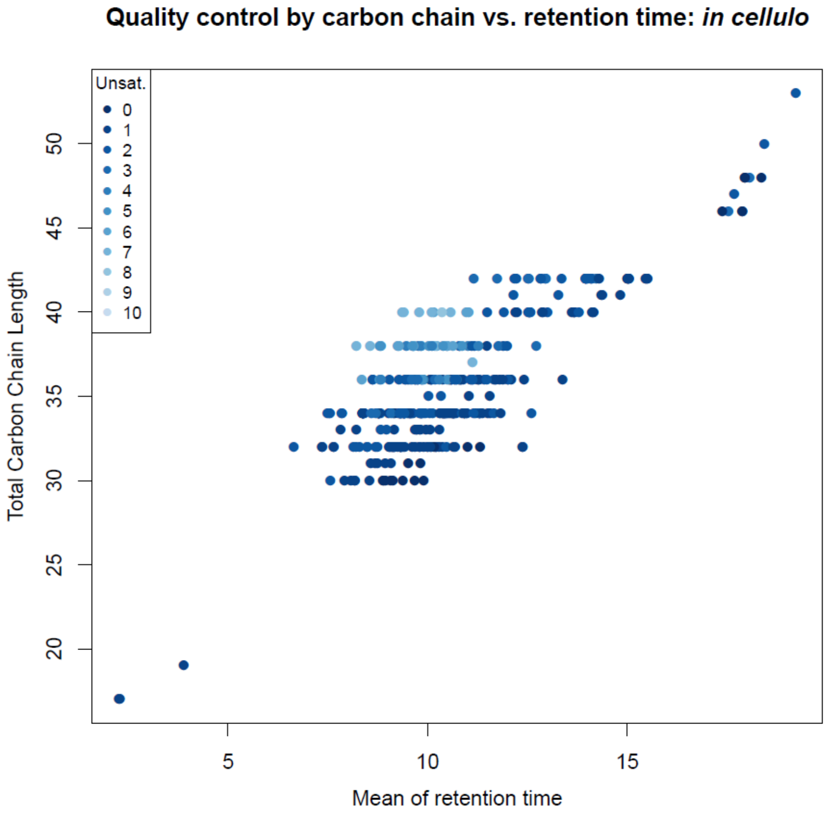

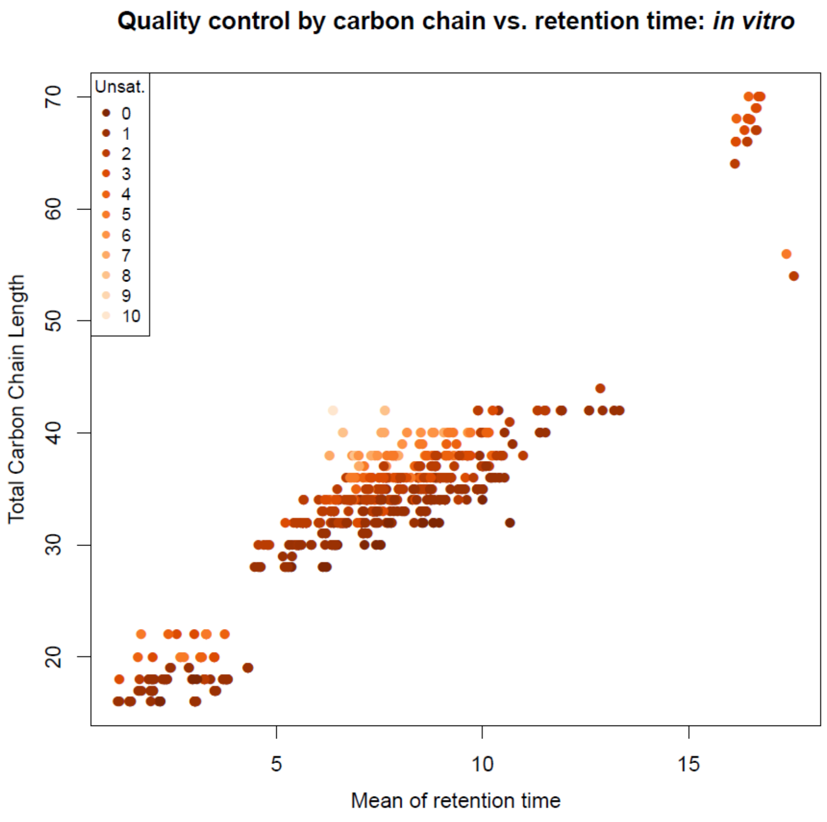


Supplementary Figure S2B: Number of LTP – lipid species combinations observed as different adducts and in different screens

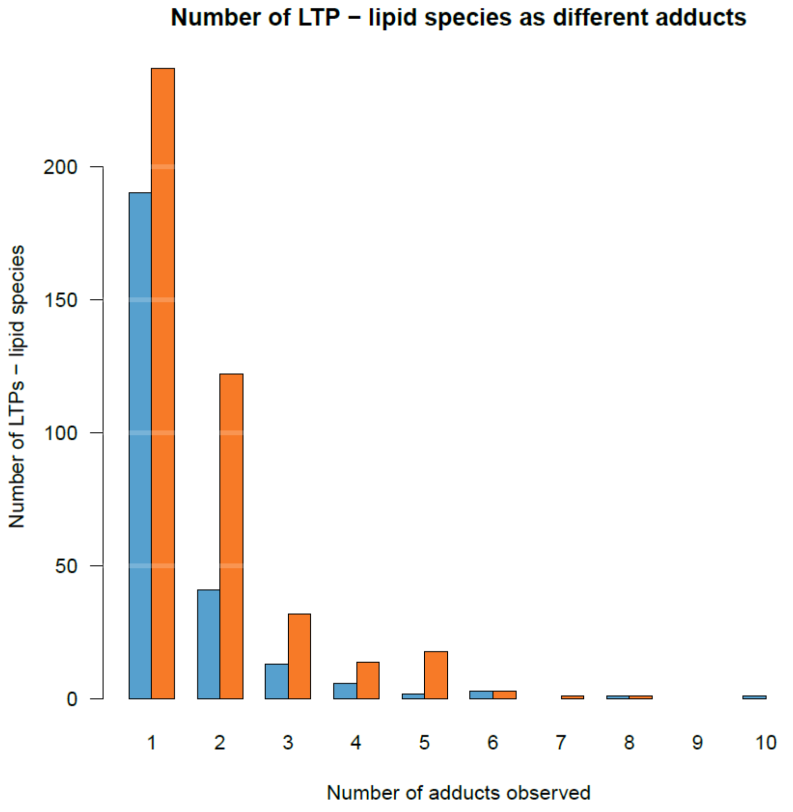

Supplementary Figure S2C: Overview of the presence of odd and even species for each of the observed headgroup categories.

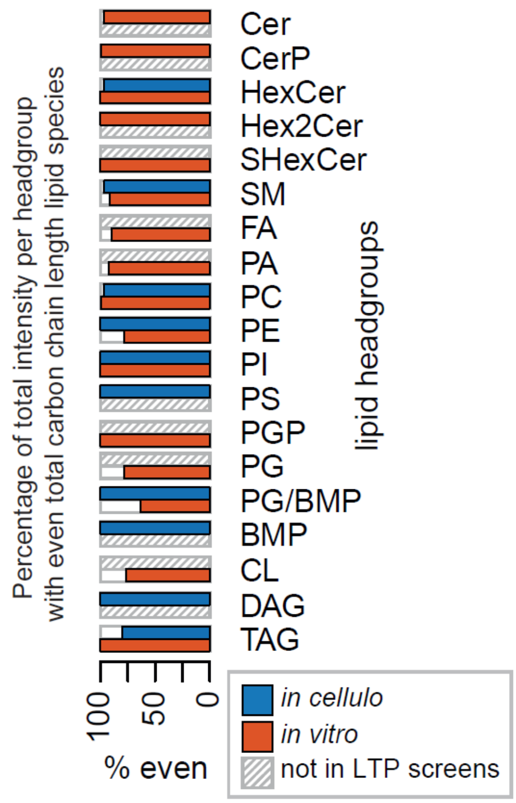


Supplementary Figure S3: Observed lipid species of phosphatidylinositol (PI) and diacylglycerol (DAG) for SEC14L2 (*in cellulo*: in filled blue) in comparison with the observed species for these lipid classes in the full HEK293 cells (empty grey boxes).

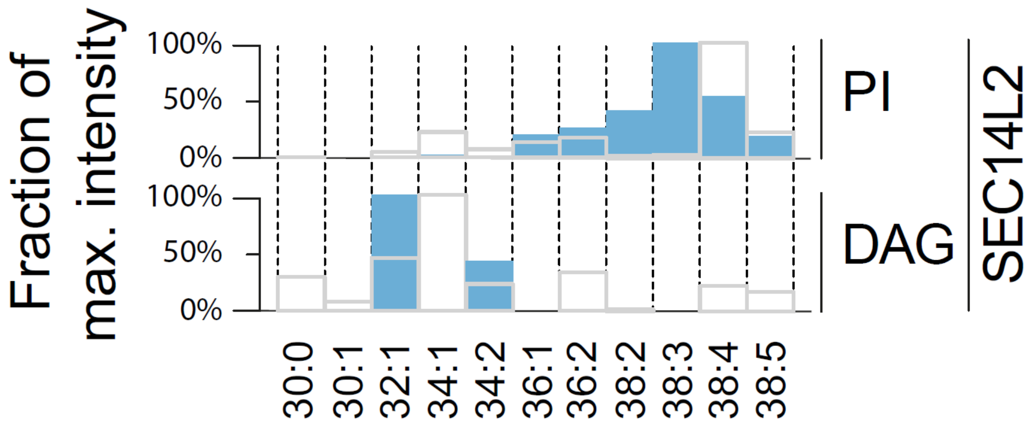

Supplementary Figure S4: Results of the seriation of the LTPs based on the location of protein domains, regions and motifs for the orderings of the main graphs and data.

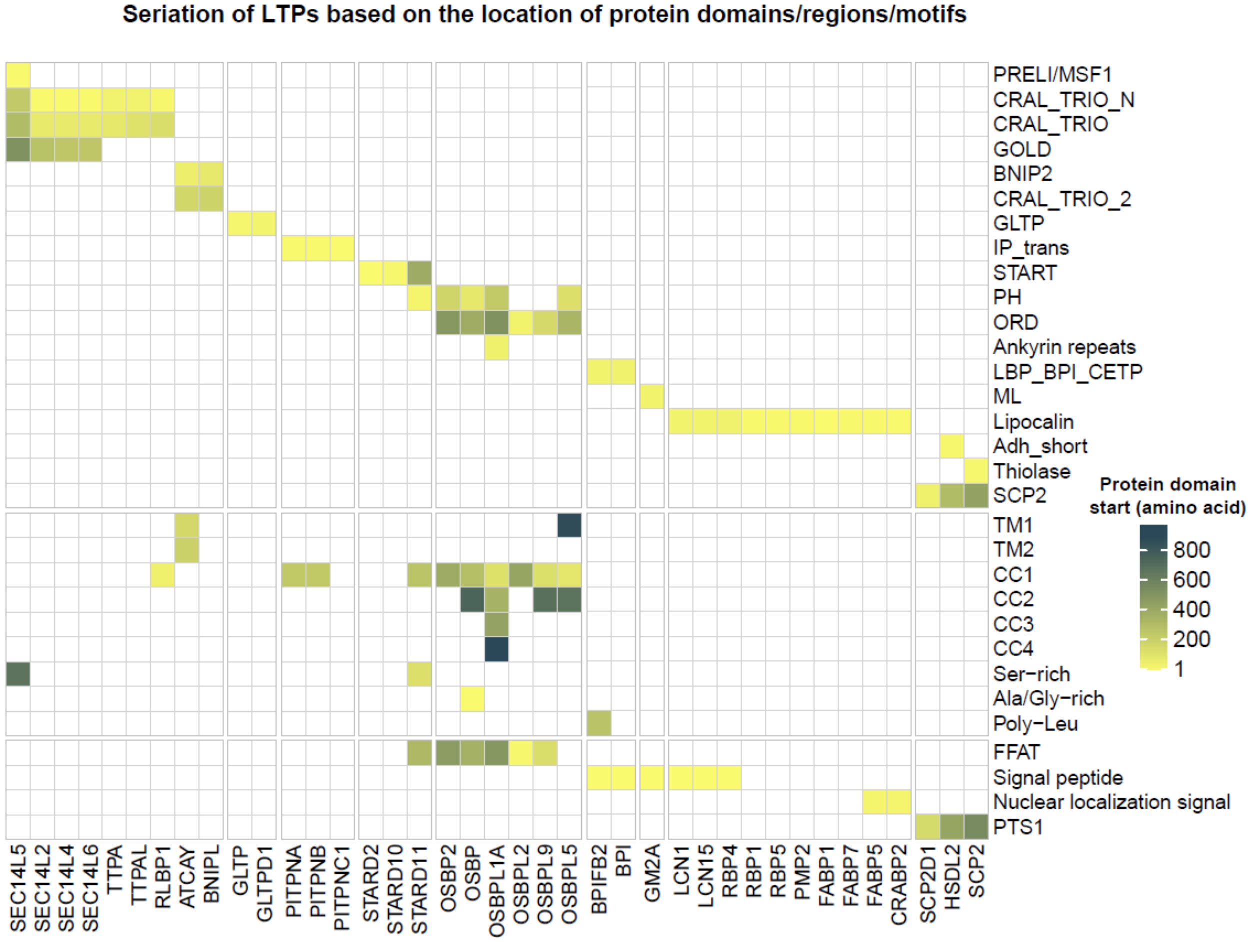

Supplementary Figure S5A: Analysis of the HSDL2 interactome and associated enrichment of its characteristics.

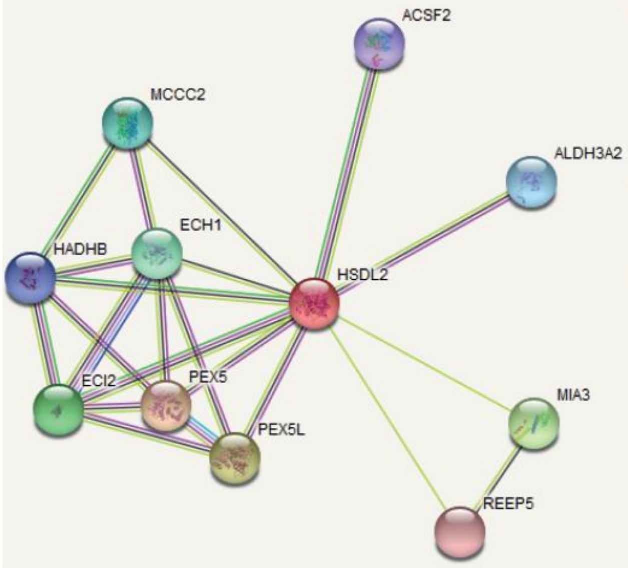


**Edges:**

Edges represent protein-protein associations

*associations are meant to be specific and meaningful, i.e. proteins jointly contribute to a shared function; this does not necessarily mean they are physically binding each other.*

| Known Interactions | Predicted Interactions | Others |
|---|---|---|
| *from curated databases* | *gene neighborhood* | *textmining* |
| *experimentally determined* | *gene fusions* | *co-expression* |
| | *gene co-occurrence* | *protein homology* |

**Your Input:**

| | |
|---|---|
| HSDL2 | *Hydroxysteroid dehydrogenase-like protein 2; Has apparently no steroid dehydrogenase activity; Belongs to the short-chain dehydrogenases/reductases (SDR) family (418 aa)* |

**Predicted Functional Partners:**

| | | Score |
|---|---|---|
| PEX5 | *Peroxisomal targeting signal 1 receptor; Binds to the C-terminal PTS1-type tripeptide peroxisomal targeting signal (SKL-type) ...* | 0.789 |
| PEX5L | *PEX5-related protein; Accessory subunit of hyperpolarization-activated cyclic nucleotide-gated (HCN) channels, regulating th...* | 0.725 |
| MIA3 | *Transport and Golgi organization protein 1 homolog; Plays a role in the transport of cargos that are too large to fit into COPII-...* | 0.644 |
| ECI2 | *Enoyl-CoA delta isomerase 2, mitochondrial; Able to isomerize both 3-cis and 3-trans double bonds into the 2-trans form in a ...* | 0.633 |
| ECH1 | *Delta(3,5)-Delta(2,4)-dienoyl-CoA isomerase, mitochondrial; Isomerization of 3-trans,5-cis-dienoyl-CoA to 2-trans,4- trans-dien...* | 0.628 |
| MCCC2 | *Methylcrotonoyl-CoA carboxylase beta chain, mitochondrial; Carboxyltransferase subunit of the 3-methylcrotonyl-CoA carbox...* | 0.590 |
| ALDH3A2 | *Fatty aldehyde dehydrogenase; Catalyzes the oxidation of long-chain aliphatic aldehydes to fatty acids. Active on a variety of ...* | 0.579 |
| HADHB | *hydroxyacyl-CoA dehydrogenase trifunctional multienzyme complex subunit beta (474 aa)* | 0.578 |
| ACSF2 | *Acyl-CoA synthetase family member 2, mitochondrial; Acyl-CoA synthases catalyze the initial reaction in fatty acid metabolis...* | 0.575 |
| REEP5 | *Receptor expression-enhancing protein 5; May promote functional cell surface expression of olfactory receptors; Belongs to t...* | 0.571 |

**Biological Process (GO)**

| GO-term | description | count in gene set | false discovery rate |
|---|---|---|---|
| GO:0046395 | carboxylic acid catabolic process | 5 of 237 | 1.23e-05 |
| GO:0032787 | monocarboxylic acid metabolic process | 6 of 477 | 1.23e-05 |
| GO:0019395 | fatty acid oxidation | 4 of 75 | 1.23e-05 |
| GO:0009062 | fatty acid catabolic process | 4 of 84 | 1.23e-05 |
| GO:0006631 | fatty acid metabolic process | 5 of 294 | 1.23e-05 |
| GO:0006625 | protein targeting to peroxisome | 4 of 68 | 1.23e-05 |
| GO:0016560 | protein import into peroxisome matrix, docking | 2 of 3 | 3.81e-05 |
| GO:0006635 | fatty acid beta-oxidation | 3 of 56 | 5.29e-05 |
| GO:0055114 | oxidation-reduction process | 5 of 923 | 0.00072 |
| GO:0046907 | intracellular transport | 5 of 1390 | 0.0038 |
| GO:0015031 | protein transport | 5 of 1391 | 0.0038 |
| GO:0034032 | purine nucleoside bisphosphate metabolic process | 2 of 123 | 0.0108 |
| GO:0033875 | ribonucleoside bisphosphate metabolic process | 2 of 123 | 0.0108 |
| GO:0006732 | coenzyme metabolic process | 2 of 297 | 0.0471 |

**Molecular Function (GO)**

| GO-term | description | count in gene set | false discovery rate |
|---|---|---|---|
| GO:0005052 | peroxisome matrix targeting signal-1 binding | 2 of 2 | 0.00012 |
| GO:0033218 | amide binding | 3 of 321 | 0.0121 |
| GO:0016874 | ligase activity | 2 of 155 | 0.0369 |
| GO:0016853 | isomerase activity | 2 of 147 | 0.0369 |
| GO:0016614 | oxidoreductase activity, acting on CH-OH group of donors | 2 of 135 | 0.0369 |

**Cellular Component (GO)**

| GO-term | description | count in gene set | false discovery rate |
|---|---|---|---|
| GO:0005777 | peroxisome | 6 of 127 | 2.80e-09 |
| GO:0044439 | peroxisomal part | 5 of 96 | 3.53e-08 |
| GO:0005782 | peroxisomal matrix | 3 of 53 | 4.58e-05 |
| GO:0005778 | peroxisomal membrane | 3 of 51 | 4.58e-05 |
| GO:0005739 | mitochondrion | 6 of 1531 | 0.00060 |
| GO:0044444 | cytoplasmic part | 11 of 9377 | 0.0022 |
| GO:0043231 | intracellular membrane-bounded organelle | 11 of 10365 | 0.0060 |
| GO:0005759 | mitochondrial matrix | 3 of 463 | 0.0113 |
| GO:0098588 | bounding membrane of organelle | 5 of 1950 | 0.0123 |
| GO:0044446 | intracellular organelle part | 10 of 8882 | 0.0123 |
| GO:0098805 | whole membrane | 4 of 1554 | 0.0297 |
| GO:0070013 | intracellular organelle lumen | 7 of 5162 | 0.0351 |
| GO:0005783 | endoplasmic reticulum | 4 of 1796 | 0.0410 |
| GO:0098827 | endoplasmic reticulum subcompartment | 3 of 1025 | 0.0480 |

Methodology:
Combination of STRING analysis of HSDL2 interactome & GO enrichment on these genes (database version: 15/06/2020).

Supplementary Figure S5B: Analysis of the peroxisomal targeting signal (PTS1) for HSDL2 and the other LTPs from the SCP2 family which mobilized lipids in our screens (based on the indicated patterns from PSORT and PROSITE).

| | **SCP2** | **HSDL2** | **SCP2D1** |
|---|---|---|---|
| C-terminal motif | AKL | ARL | AKF |
| [A/C/S]–[H/K/R]–L (PSORT) | ok | ok | not ok |
| [A/C/G/N/S/T]–[H/K/R]–[A/F/I/L/M/V/Y] (PROSITE) | ok | ok | ok |
