## Supplementatary methods for "A system-wide analysis of lipid transfer proteins delineates lipid mobility in human cells"

**Material and methods**

**Selection and cloning of tagged recombinant human LTPs**

We selected 125 human LTPs from literature and from protein (domain) databases (UniProtKB, SMART and InterPro) (Chiapparino et al., 2016, Prog Lipid Res). Of these, 101 LTPs were cloned in frame with an N-terminal His6-HA-StrepII-tag in a pcDNA5/FRT/TO vector for expression in HEK293 cells (gift from the Beck laboratory, based on addgene construct 2133). Coding sequences of the full-length LTPs were obtained from commercial cDNA libraries (the Human Universal cDNA library of BioCat, Open Biosystems, Harvard Medical School), as well as from in house generated cDNA libraries. For the generation of cDNA libraries, we extracted the mRNA from HeLa, HEK293, MCF7 cells and mesenchymal stem cells using the RNeasy mini kit (Qiagen) and performed SuperScript™III RT-PCR. The primers used for the amplification of the cDNAs are described in Supplementary Data S1C. Each primer extended the two edges of the template cDNA with SfiI restrictions sites, compatible with SfiI restriction sites present on the modified pcDNA/FRT/TO plasmids in frame with the tandem affinity purification cassette. The inserted cDNAs were sequenced with both forward and reverse sequencing primers.

Using a modified version of pETM11-SUMO3 with SfiI enzyme restriction sites (provided by the EMBL Protein Expression and Purification Core Facility), an LTP-library was generated for the expression of LTPs in *E. coli*. The pETM11 vector has an N-terminal His6-TEV-SUMO3 tag and a Kanamycin resistance gene allowing antibiotic selection. The expression of genes for the pETM11 vector was under control of the Lac operon for induction of protein expression in *E.* coli. During the cloning, the cut fragments by SfiI were treated with alkaline phosphatase, and the cut inserts were separated on and purified from a 0.8% agarose gel, while the digested vectors were similarly separated from the undigested vector and possible contaminants on a 1% agarose gel. For transformation, we employed XL1-Blue Competent *E. coli*. The strains were grown overnight in Kanamycin plates at 37°C. Colony PCR was used to evaluate the presence of inserts, and the gene sequence was checked by sequencing the first 150-200 nucleotides after the first SfiI site.

**Generation of the stable human cell lines (*in cellulo* approach)**

HEK293 cells were maintained in DMEM, supplemented with 10% (v/v) FBS, 1% L-glutamine, and kept under selection in the presence of 15 µg/ml blasticidin and 100 µg/ml Zeocin before transfection. To create stable inducible human cell lines for LTPs, Flp-In T-Rex-293 cells were co-transfected with the plasmid coding for the tagged LTP and the pOG44 plasmid encoding the Flp recombinase (Invitrogen). Positive clones were selected by adding 100 µg/ml hygromicin B and 15 µg/ml blasticidin on the day after transfection. The generation of the stable cell lines lasted each 3-4 weeks, and we generated 40 cell lines in parallel per month. In this timeframe, we expanded the cell lines in the presence of 1 µg/ml tetracycline (without other antibiotics) up to 40 p150 Thermo Scientific™ Nunc™ Cell Culture Dishes, whereafter the cells were grown further until 95% confluence (3-4 days), harvested, pelleted (yielding 2-3 ml pellet), and stored at -80°C for later use. Protein expression was evaluated by Western blot with alpha-HA antibody (Supplementary Table S3), and all cell lines were tested for the presence of mycoplasma.

**Protein expression and LTP-complex purifications from cell lysates (*in cellulo* approach)**

Cell lysis was performed by resuspending cell pellets in lysis buffer (50 mM Tris-HCl, 250 mM NaCl, 0.5 mM DTT; 2 µM avidin, protease inhibitors cocktail (Roche) and DNAse (Roche), at pH 7.4) and leaving on ice for 20 minutes. In the case of membrane proteins and proteins closely associated to membranes, 0.5% NP40 was added to the lysis buffer to facilitate solubilization (Supplementary Table S3).

The final cell extract was obtained by centrifugation for 20 min at 16,000 g at 4°C in a bench top centrifuge (Eppendorf 5427R), followed by centrifugation of the previous supernatant at 49,000 rpm, using a TLA100.4 rotor. Protein-lipid complexes were isolated via the Strep-II tag, at room temperature and eluted with 5 mM biotin. The eluted complexes were centrifuged at 16,000 g for at least 5 minutes and fractionated on a Superdex 200 SEC column (Invitrogen). After elution, protein fractions were kept separated. A part of them was used for sodium dodecyl sulfate–polyacrylamide gel electrophoresis (SDS-PAGE) on pre-cast 4-12% gradient gels (Life technologies), while the major part (100 µl) underwent lipid extraction. We performed High Performance Thin Layer Chromatography (HPTLC) in the case of the likely presence of ligands difficult to detect with mass spectrometry because of their chemical nature, such as cholesterol and cholesterol esters.

**Protein expression in *E. coli* (*in vitro* approach)**

The His-SUMO3 tagged proteins were expressed in BL21 Star *E. coli* cells (Invitrogen) grown in ZYM supplemented medium at 37°C until an OD600 of 1.5. Immediately after, the temperature was dropped to 18°C for the following 14h of protein overexpression. Cells were harvested at 4.000 rpm in JLA8100 rotor, resuspended and washed with PBS, centrifuged again in 50 mL falcon tubes in a Centrifuge 5810 and snap frozen with liquid nitrogen. The *E. coli* cell pellets were stored at -80°C until the day of purification.

**Liposome preparation (*in vitro* approach)**

Dioleoyl phosphatidylcholine (DOPC), porcine brain polar lipid extracts and bovine liver total lipid extract were obtained from Avanti Polar Lipids. DOPC was used as a carrier lipid to help the formation of liposomes. A mixture of 1:1 molar ratio of DOPC and liver lipid extract and another mixture of 1:1 molar ratio of DOPC and brain lipid extract were prepared. Both solutions had a total amount of lipids of 3 mg. The lipid mixtures were dried into thin lipid films in glass-vials by a flow of argon gas followed by vacuum for at least 30 minutes. One milliliter of rehydration buffer (50 mM HEPES pH 7.4, 250 mM NaCl, 750 mM sucrose) was added to each lipid film in order to form liposomes to a total lipid concentration of 3 mg/ml. Vials were shaken vigorously at 62°C for at least 1h, and liposomes were subsequently downsized to 400 nm by filtering 21 times through Whatman Nuclepore Track-Etched Membranes on a mini-extruder (Avanti Polar Lipids).

**Incubation of LTPs with mammalian tissue derived liposomes (*in vitro* approach)**

Cell pellets were thawed on ice and resuspended in the lysis buffer (50 mM Tris pH 7.5, 250 mM NaCl, 20 mM imidazole, 0.5 mM DTT, protease inhibitors: Pefabloc 1 mM (Sigma), Bestatin 3 μM (Sigma), Aprotinin 300 nM (Sigma), E64 1 μM (Sigma), Pepstatin A 1.5 μM (Sigma), Leupeptin 1 μM (Sigma). Complete cell lysis was achieved by pressurization in Avestin Emulsiflex C3. Cell membranes and aggregated proteins were eliminated by centrifugation at 21,000 rpm for 1h in the JLA25.50 rotor. The soluble fraction was incubated with bovine liver liposomes (1:250) and porcine brain liposomes (1:250) (see next paragraph), and incubated for 1h at 4°C in in a roller mixer for falcon tubes.

Afterwards, the liposomes were separated from the lysate by centrifugation at 21,000 rpm for 1 h in the JLA25.50 rotor. Cleared lysate was incubated with Ni-NTA agarose resins (Qiagen) in order to capture the protein-lipid complexes. The affinity columns were washed with lysis buffer without protease inhibitor cocktail and cleaved overnight with TEV or SenP2 proteases. The eluted protein-complexes were loaded on a Tricorn5/150 Superdex 200 self-packed size-exclusion chromatography colum. After this, the fractions were collected to undergo the same further analysis steps as for the *in cellulo* approach.

**Lipid extraction**

For HPTLC analysis, lipids were extracted from samples by sequential addition of 3.75 volume chloroform:methanol (1:2 v/v), 1.25 volume chloroform, and 1.25 volume 0.5% acetic acid in 500mM NaCl followed by 30 seconds of vortexing after each step. After centrifugation at 1,200 rpm for 10 min, the bottom layer was dried under vacuum at RT in the dark and resuspended in 25 µl of a solution of chloroform:methanol (2:1).

Samples for lipid-MS analysis were prepared in the same way as described above, with a few modifications as follows. After lipid extractions, the extracted lipids were dried and resuspended in 65 µl methanol in the same glass vial. The samples were later split in two fractions and transferred into a 96-well plate (twin.tec PCR Plate 96) to be analyzed in in both positive and negative ionization mode.

**Lipid analysis by high-performance thin layer chromatography**

Extracted and resuspended lipids were sprayed (in 25 µl) on HPTLC silica gel 60 plates (10 cm x 10 cm; Merck) in 3 mm bands using the TLC-sampler 4 (Camag). The plates were sequentially developed with a solvent system of (1) chloroform:methanol:water:acetate 65:25:4:1, and (2) cyclohexane:ethylacetate (1:1) for the analysis of polar and neutral lipids (modified from Weerheim et al., 2002, Analytical Biochemistry). The plates were dried under vacuum and dripped in 10% (w/v) CuSO_4_ in 8% (v/v) aqueous phosphoric acid, charred at 145°C for 4.5 min (Churchward et al., 2008, J Chem Biol), and scanned for fluorescence detection using Pharos FX Plus molecular imager (Bio-Rad; 488 nm) (488 nm excitation and 530 nm emission wavelengths). Lipids applied as standards were detected with a sensitivity of <1 ng. Images were analyzed and processed using ImageJ.

**Lipid analysis by liquid chromatography followed by tandem mass spectrometry**

Lipids (65 µl) were separated on an Agilent 1260 HPLC system consisting of a degasser, binary pump, and autosampler directly coupled to a Q-Exactive Plus (Thermo), equipped with a heated ESI source. The column was a Kinetex 30 x 2.1 mm, 2.6 µm, C18, 100 Å (Phenomenex). A binary solvent system was used in order to separate lipids. The mobile phase A consisted of H_2_O:acetonitrile (60:40), 10 mM ammonium formate, and 0.1% formic acid, while the mobile phase B consisted of isopropanol:acetonitrile (90:10), 10 mM ammonium formate, and 0.1% formic acid. The separation started at 80% buffer A and 20% buffer B. In a 3 min gradient, buffer B was increased from 20% to 50%, followed by a 10 min gradient from 50% to 70% buffer B. Finally, a 5.4 min gradient was applied to increase buffer B from 70% to 97%. The column was subsequently washed for 2.1 min with 97% buffer B and equilibrated for 3.6 min with 20% buffer B. The flow rate was 500 μl/min. The entire run was 24 min long. The effluent was directly introduced into the ESI source of the MS, and analyzed each time for each paired fraction in either positive or negative ionization mode. The ESI source ion spray voltage was set to 1.7 kV. The mass spectrometer was operated in the mass range from 250 to 1600 m/z. Charge state screening was not enabled. The ten most abundant peaks (TOP 10) were selected and fragmented in MS2 by HCD. The normalized collision energy (NCE) was 30. MS1 resolution was set to 70,000 at 200 m/z, while for the MS2 it was 17,500 at 200 m/z.

**Preparation of lipid standards**

To ensure maximal accuracy and depth of identification to the lipid species level, lipid standards for each lipid subclass were analyzed by LC-MS/MS. This allowed to define fragmentation pattern and retention time rules for each lipid subclass. Lipid standards were dissolved in CHCl_3_MeOH (2:1 v/v) at concentrations ranging from 1 to 10 mg/ml and were stored at -20°C. The targeted lipid subclasses were “PC”, “PC-O”, “LPC”, “PE”, “PE-O”, “LPE”, “LPE-O”, “PS”, “PI”, “PA”, “BMP”, “PG”, “LPG”, “PGP”, “CL”, “DAG”, “TAG”, “FA”, “Vitamin A”, “Vitamin E”, “sphingosine”, “sphingosine-1-phosphate”, dCerP”, “dCer”, “DHCer”, “tCer”, “dSM”, “DHSM”, “tSM”, and the same d, DH and t subclasses for “HexCer” and the related “SHexCer” and “Hex2Cer” (Avanti Polar Lipids and Sigma).

**Formulation of lipid identification rule-sets**

The lipid subclass fragmentation and elution rules derived from our standards (see previous paragraph) were confirmed by spectra in the literature and public databases, such as METLIN (Guijas et al., 2018, Anal. Chem.). The subclass rulesets are available in supplementary tables S2A&B, and the rules defined for the subclasses were automatically applied by the lipyd software and expanded to all theoretically possible lipid species to identify these (<https://github.com/saezlab/lipyd>).

**Lipid identification and filtering**

The raw MS data were converted to mzML format with MS Convert from the ProteoWizard package (version 3.0.679). PEAKS Studio 7.0 (Bioinformatics Solutions Inc.) was used for peak picking, feature detection and alignment. Features coming from each sample were grouped into a control group (buffer only and fractions before and after the protein peak) and a sample group (containing the fractions where the protein eluted). A first quality control accounted for the following criteria: quality (peak shape score from the PEAKS software) ≥0.2, (real sample/control) ratio >2 (performed for each protein sample fraction versus each control fraction), minimum peak intensity of 10000, only single charge states, and fitting with the retention time of lipids (above 1 minute). Finally, we considered the profile of the protein abundances over the different SEC fractions, and only selected the potential lipid ions with an elution profile similar to the protein profile. The lipid species were computationally identified based on the above explained fragmentation rule sets. Finally, specimens that matched the protein SEC elution profile and exceeded in intensity 100,000 and 30,000, for positive and negative ion mode, respectively, underwent a three-person manual curation that approved the final identification, using the SwissLipid database, also searching confirmation by taking the associated retention times into account. This manual curation also eliminated identifications that corresponded to the used detergent (NP40) in the lysis of some of the samples (Supplementary Table S3: samples lysed with this detergent), and we also removed any lipid identifications with ambiguity in the observed headgroups (e.g. SM vs. PC).

**MS-intensities normalization**

The MS-intensities from different screens and different lipidomics approaches were always analyzed and visualized independently, and they were normalized for representation in the (circular) heatmaps to facilitate comparisons. The intensities were first log10 transformed per mode and screen, and then min-max normalized. The min-max normalization used the minimum and maximum of the whole dataset within the particular screen and mode. These steps were followed by a scaling where maximum values correspond to 10 and minimum values to 1, while absences correspond to 0. The scaling was realized by multiplication of normalized values by 9 followed by addition of 1. After this, the results for both ion modes were compared and the maximal value was retained from either of the ion modes.

For comparisons not visualized in heatmaps and where normalization is unwanted, MS-intensity sums and percentages were used, all based on linear addition of MS-intensities without transformation. Percentages (such as in the bar charts in Fig. 2b-c) give the percentage of the considered condition in comparison with the combined intensities of the total screen (inside the particular reference frame), while percentages that indicate the fraction of the maximal intensities (such as in the bar charts in Fig. 3c & 6a) show analogously the percentage of the considered condition in comparison with the maximum intensity of the specific row.

**Protein-domain-based ordering of LTPs, and association with lipid sub-classes**

The LTPs along the x-axis of Figure 3a were ordered by seriation based on the presence of all protein domains/motifs and their location within the proteins. The result of the underlying domain/motif-based seriation is visualized in Supplementary Figure S4. The protein domains/motifs and their intra-protein locations correspond to all domains and motifs that were available for these LTPs in June 2020 in either PFAM or InterPro (El-Gebali et al., 2019, Nucleic Acids Research; Blum et al., 2020, Nucleic Acids Research). The employed seriation was according to the travelling salesman algorithm, as described in (Hahsler et al., 2008, Journal of Statistical Software).

**Comparison of lipid co-mobilization with co-regulation and co-localization**

For the validation of the observations in the screens and the discovery of biological relationships, we integrated external data from different external data sources (see further) to compare the pairs of lipids observed with the same LTPs in our data (co-mobilized lipid pairs) with their co-regulation (Fig. 5a), and co-localization within tissues (Fig. 5b) and within cells (Fig. 5c) in the external data. The next paragraph describes the shared methodological and visualization elements of these analyses, while the paragraph after describes the unique aspects of each of these analyses.

In each analysis, we extracted the data from the external data source for the observed co-mobilized lipid pairs of the here-described screens and compared their distribution with the distributions of two broader sets of lipid pairs in the external data source. One broader set of lipid pairs consisted of all theoretically possible pairwise combinations of the observed lipids, instead of only the observed combinations, while the other broader set of lipids consisted of all lipid pairs in the external data, thus also including unobserved lipids. The filled coloured distributions correspond in all panels of figure 5 to the results for the co-mobilized lipid pairs, while the coloured contour distributions correspond to the results for the theoretically possible combinations, and the grey contour distributions correspond to the results for all lipid pairs in the external data. The long vertical lines correspond to the medians of these distributions, respectively in black, blue/orange and grey. The calculation of the densities of these distributions used Gaussian kernels with bandwidth selection according to (Sheather & Jones, 1991), with an overall density of 1 in all cases corresponding to the uniform distributions. Moreover, within each analysis, all distributions have the same surface area to facilitate comparisons. All analyses were performed on the aggregate of all lipid pairs (“all” in Fig. 5), but the pairs were also split up in different subtypes of similarity to avoid domination of comparisons by closely related lipid species. The “species” subtype of lipid pairs refers to pairs that only differ in their fatty acyl chains; the “sub-class” subtype refers to pairs that differ in their linkage, for example ester-linked lipid versus ether-linked lipids or d versus DH sphingolipids; and the “class” subtype includes pairs that differ in their headgroup, for example PE and PC. The significance of the shift of the observed distribution of lipid pairs versus the external set of lipid pairs was each time determined by Fisher’s exact tests (Fisher, 1934, Statistical Methods for Research Workers, 5^th^ ed.). The median of the expected distribution was used to split both the expected and observed data, and then the observed distribution was compared with the expected distribution. Significant results were indicated with a star in the figure 5, and detailed results of the comparisons can be found in supplementary table S4.

The data used to study the co-regulation of co-mobilized lipids (Fig. 5a) was derived from (Köberlin et al., 2015, Cell). Co-regulations in these data represent Pearson’s linear correlations in the log2-transformed changes in abundance of the lipids of the lipid pairs upon cellular perturbations. For the analysis of the co-localization in tissues (Fig. 5b), we calculated the Manders’ co-occurrence of all pairs of lipid-like molecules (as defined by ClassyFire (Feunang et al., 2016, J Cheminform)) in the mass spectrometry based imaging metabolomics database METASPACE (Manders et al., 1993, J Microsc Oxford; Alexandrov et al., 2019, bioRxiv). All datasets from human tissues in this database were selected up to May 2018, if there were at least 50 of the co-transported lipid species present per dataset (below 5% FDR). The background observations were removed by selecting for each dataset a metabolite with uniform distribution across the tissue, followed by morphological transformation of the corresponding image with grayscale erosion and then grayscale dilation, to define a mask representing the tissue region. We subsequently also discarded datasets where the average intensity of the background was higher than the average intensity of the tissue region. The co-occurrences of the lipid pairs were calculated for all pixels corresponding to tissue for each of the datasets. The use of the Manders’ overlap coefficient allowed to eliminate the influence on calculation of the co-localizations of the absence of both queried lipids in pixels. For analysis of the sub-cellular co-occurrence of the co-mobilized lipids (Fig. 5C), we also employed the Manders’ overlap coefficient, but on a dataset containing the lipidomes of organelles purified with antibodies (article in preparation by the Maeda lab, Denmark).

**Data analysis scripts availability and programming context**

The in-house developed library for lipid identification and matching of lipid elution profiles with protein protein elution profile is called Lipyd and is accessible via <https://github.com/saezlab/lipyd>. The R scripts for the data clean-up, and data analyses and visualizations are available via <https://github.com/krtiteca/ScriptsAssociatedWithLTPArticle>. For these analyses and visualization, we consequently used R version 3.5.0. For the visualization of beanplots, we employed the R package by (Kampstra, 2008, Journal of Statistical Software), and for the circular visualization and heatmaps, we employed the packages circlize and ComplexHeatmap (Gu et al., 2014, Bioinformatics; Gu et al., 2016, Bioinformatics).

**Systems preparation for molecular dynamics simulation**. We recently reported simulations of START apo and holo in presence of complex lipid bilayers mimicking the ER and Golgi membranes ^1^. In this work we observed the insertion of one tail of a POPC (1-palmitoyl-2-oleoyl-phosphatidylcholine) lipid in the START hydrophobic cavity. We extracted structures of the apo and holo START domains together with POPC lipid from the simulation trajectories. We then performed a single point energy calculation on the extracted complexes using the CHARMM-GUI PDB Reader & Manipulator ^2,3^. This step ensured that the atomic coordinates were successfully defined and that the structure was ready to use in CHARMM-GUI Solution Builder tool ^4,5^. The complexes were solvated in a cubic box of size 90 × 90 × 90 nm^3^ using explicit TIP3P water molecules and neutralized with three potassium ions. To ensure that the final systems had no steric clashes or inappropriate geometry, an energy minimization was performed prior to starting a 400 ns-long dynamics simulation.

**Simulations protocol.** Our MD simulations were performed using NAMD (v 2.13) ^6^ with the CHARMM36 force field ^7^ and its CHARMM-WYF extension for the treatment of aromatics-choline interactions ^8,9^. Before the MD simulations were carried out, each system was subjected to energy minimization with conjugate gradients (10000 steps). After a short equilibration run (5 ns) in the NVT ensemble, the production runs were performed for at least 400 ns with an integration step of 2 fs in the NPT ensemble. Temperature and pressure were set at 310 K and 1 bar, respectively. Langevin dynamics with a temperature damping coefficient of 1.0 and the Langevin piston method with an oscillation period of 50 fs and a damping timescale of 25 fs were used to control the temperature and pressure, respectively. Lengths of all bonds between hydrogen and heavy atoms, including those in water molecules to keep water molecules rigid, were controlled via the SHAKE algorithm ^10^. Electrostatic potentials were calculated using the Particle Mesh Ewald (PME) method ^11^. A Lennard-Jones switching function over the range of 10-12 Å was used for van der Waals interactions. Within simulation time, the second tail of the POPC lipid entered the cavity.

**Visualization.** Image generation was done by PyMOL ^12^.
