## Supplementary Data S1A,B for "A system-wide analysis of lipid transfer proteins delineates lipid mobility in human cells"

**Supplementary Data S1A:
Purification and characterization of LTPs-lipid complexes *in cellulo* (with SEC, SDS-PAGE and HPTLC data)**

LTP-lipid complexes were isolated from the respective cell lines and fractionated on

Superdex 200 SEC column. Chromatogram (A215), SDS-PAGE, and HPTLC analysis of

GPL, SL and sterols of LTPs isolated from the respective human cell lines. On top of each

chromatogram, the resolution of the column is displayed. **a,** OSBP domain proteins**. b,**

START domain proteins. **c,** ML domain protein. **d,** CRAL-TRIO domain proteins. **e**, SDS-PAGE and HTPLC are shown for the START domain protein, PCTP (synonym of STARD2) and for the GLTP domain protein, GLTP. These two proteins were fractionated on a different SEC column.
*The indicated region of the HPTLC is displayed with enhanced contrast below or next the full HPTLC image. PIPs, phosphatidylinositol phosphate; PC, phosphatidylcholine; PI, phosphatidylinositol; PS, phosphatidylserine; PA, phosphatidic acid; CA, cardiolipin; PE, phosphatidylethanolamine; Chol, cholesterol; CO, cholesteryl oleate; SphP, sphingosine-1-phosphate; SM, sphingomyelin; CerP, ceramide-1-phosphate; Cer, ceramide; S, standard

containing GPL, cholesterol and CO; S2, standard containing SL (see CERT binding profile,

panel **d**). S3 contains Erg (ergosterol) instead of cholesterol and does not contain CO (only used for experiments in panel **e**). E, elution

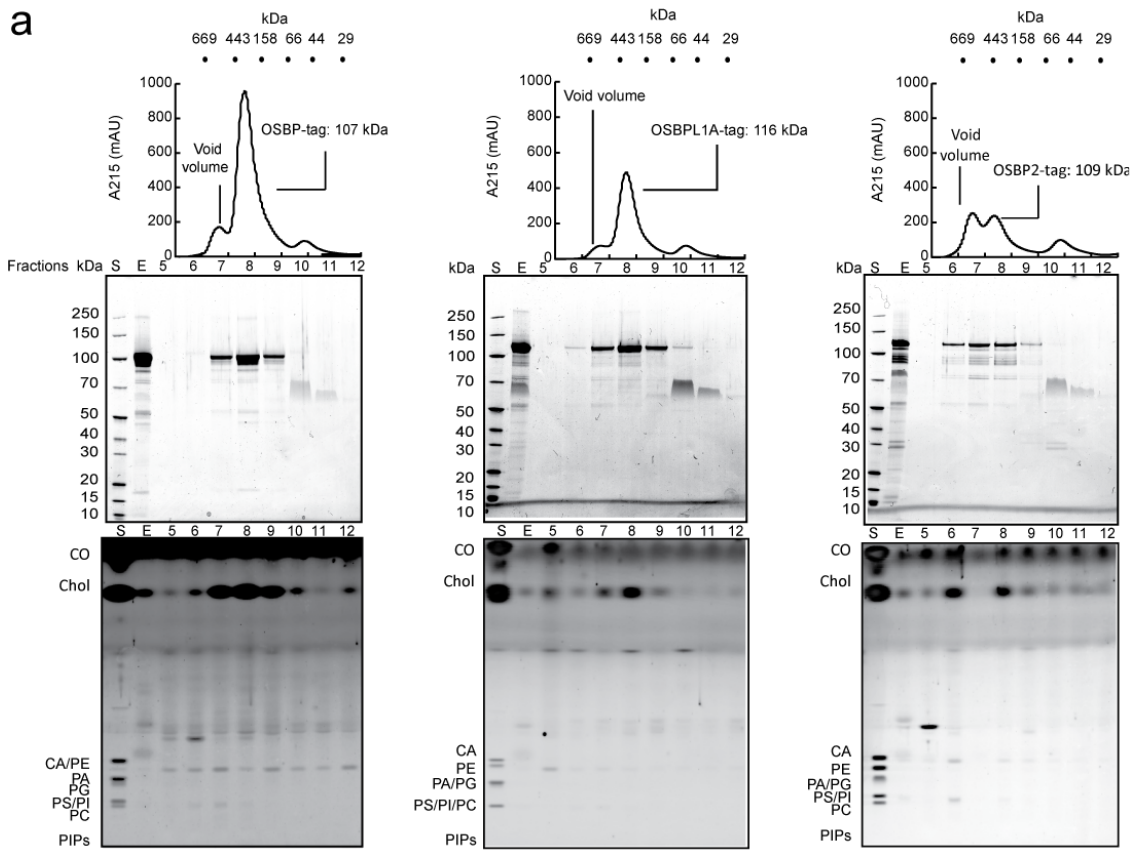

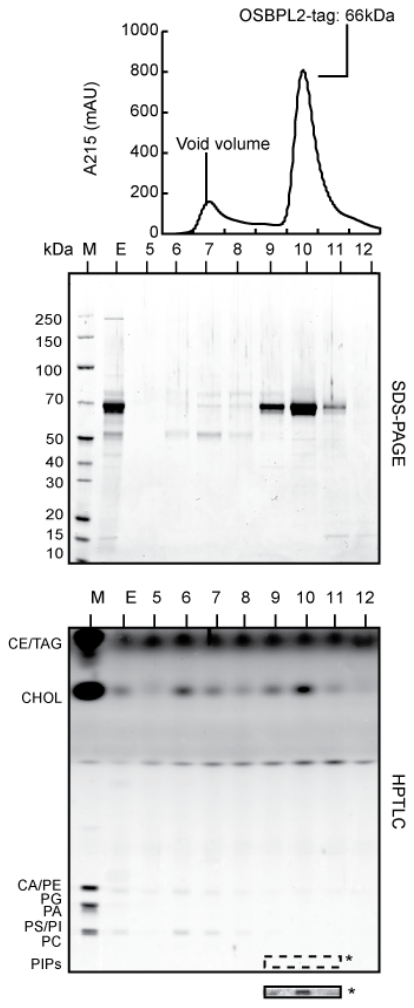

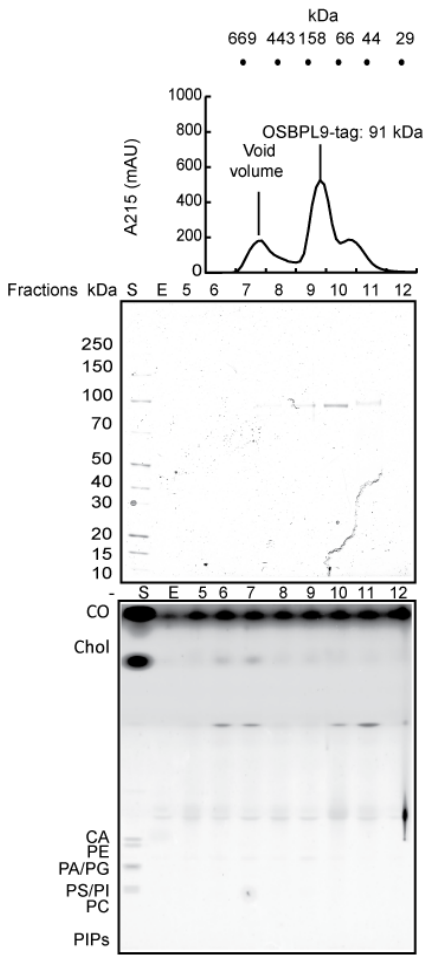

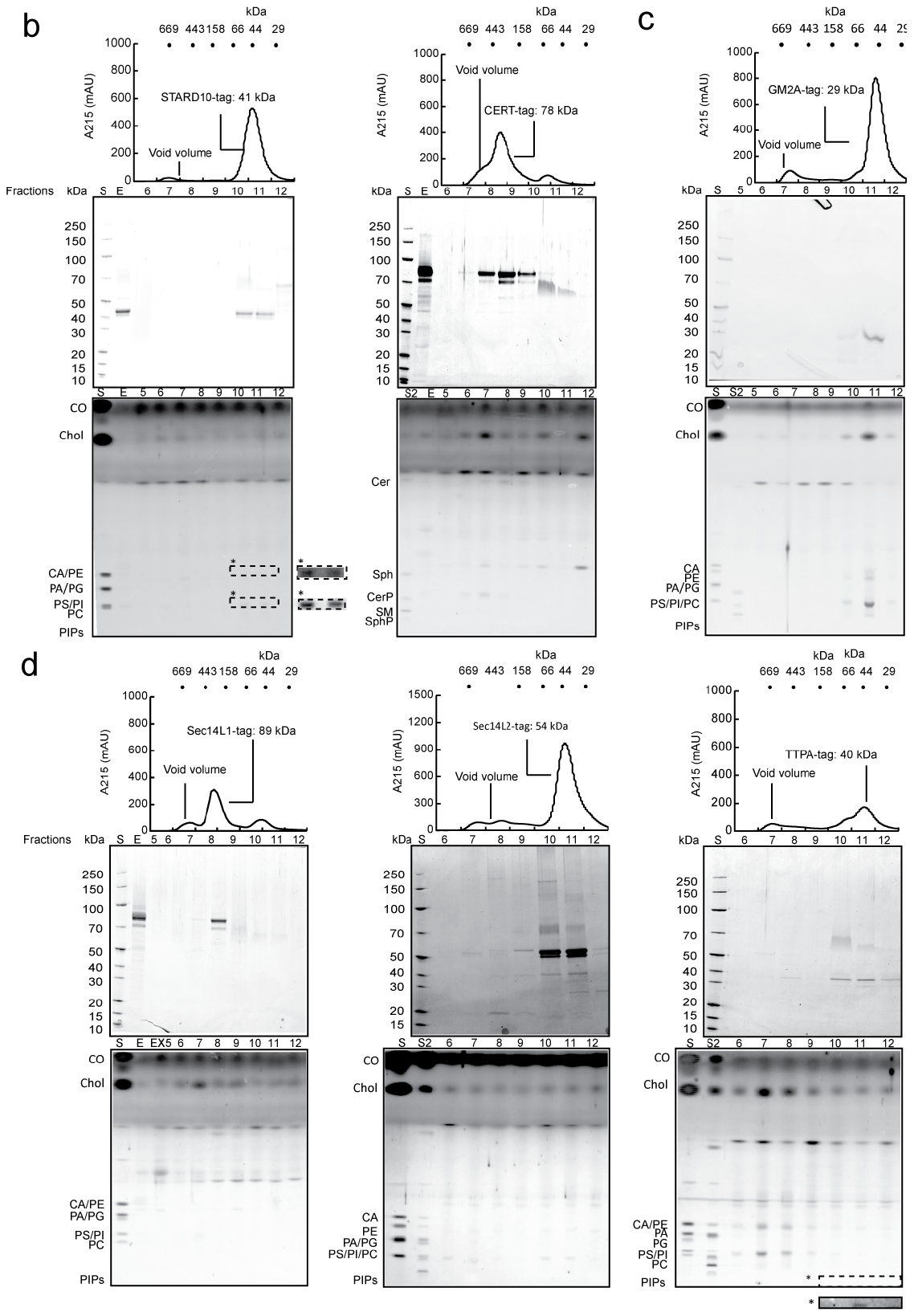

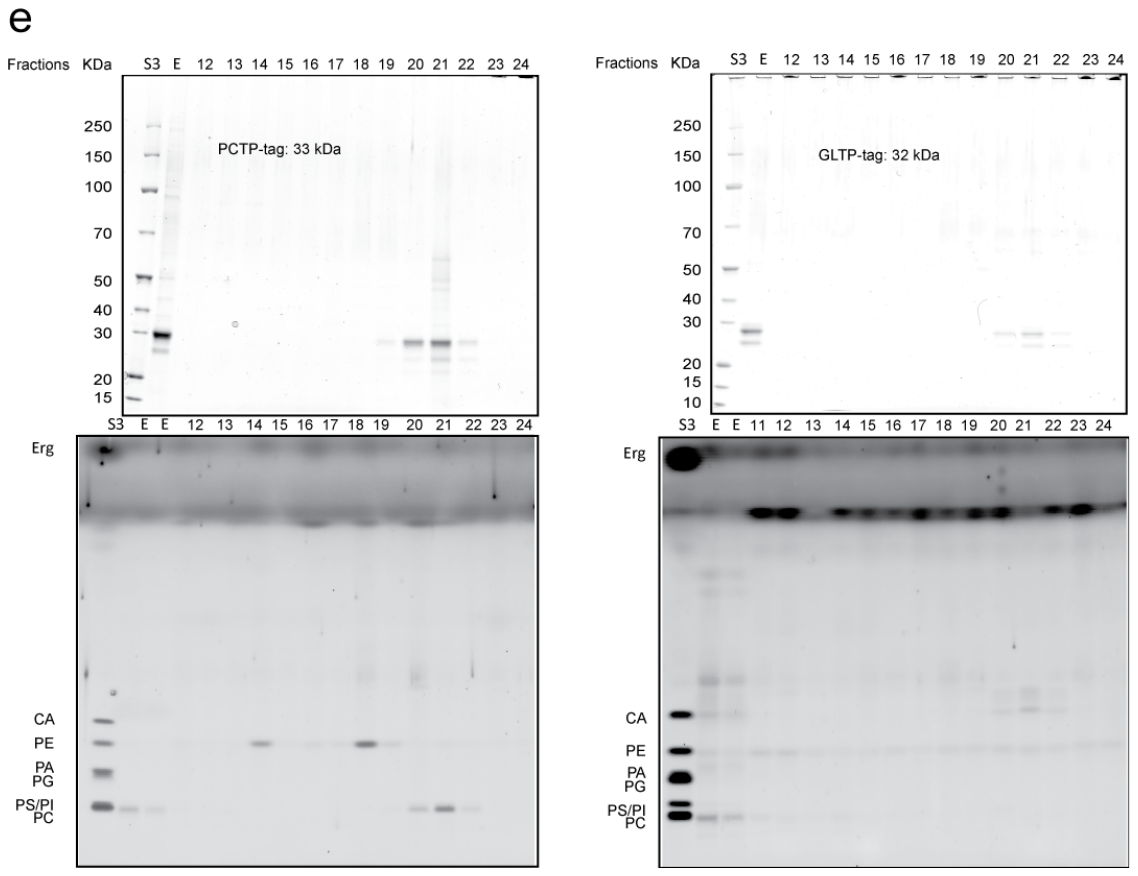

(STARD2)

**Purification and characterization of LTPs-lipid complexes *in cellulo* (with SEC and SDS-PAGE data)**

Chromatogram (A215) and SDS-PAGE of LTPs isolated from the respective human cell

lines. On top of each chromatogram, the resolution of the column is displayed. **a,** OSBP

domain proteins. **b,** lipocalin domain proteins. **c,** CRAL-TRIO domain proteins. **d,** SCP2

domain protein; **e,** PITP domain proteins. **f,** GLTP domain protein. **g,** NPC1 NTD domain

protein. **h,** START domain proteins. **i,** LBP_BPI_CETP domain proteins. PIPs,

phosphatidylinositol phosphate; PC, phosphatidylcholine; PI, phosphatidylinositol; PS,

phosphatidylserine; PA, phosphatidic acid; CA, cardiolipin; PE, phosphatidylethanolamine;

Chol, cholesterol; CO, cholesteryl oleate; SphP, sphingosine-1-phosphate; SM,

sphingomyelin; CerP, ceramide-1-phosphate; Cer, ceramide; E, elution.

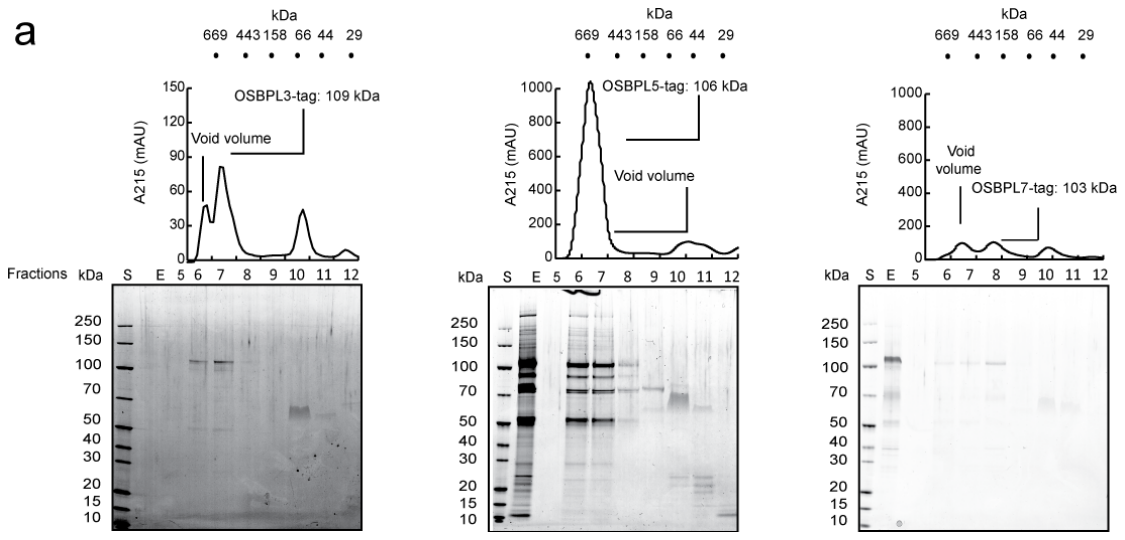

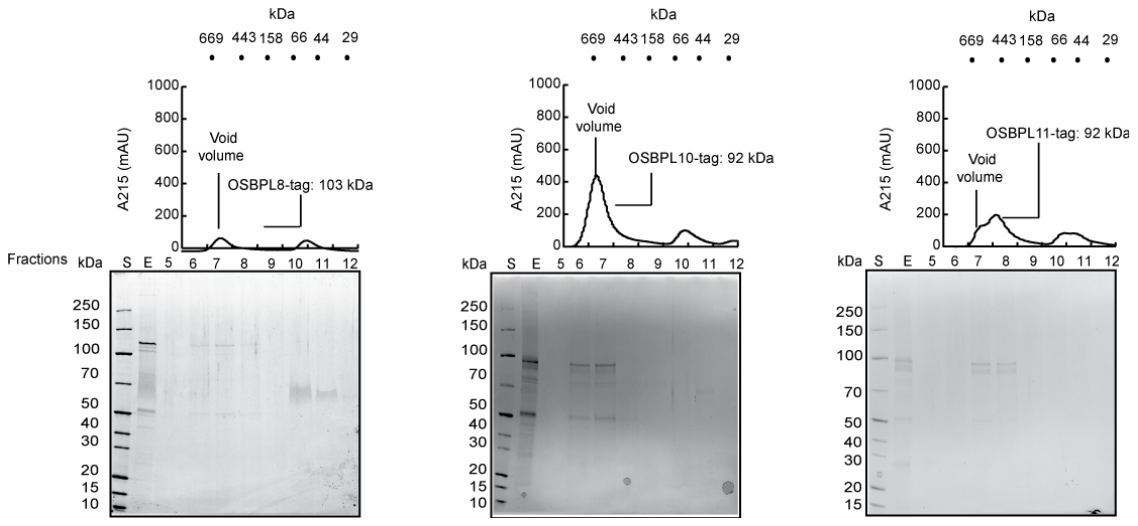

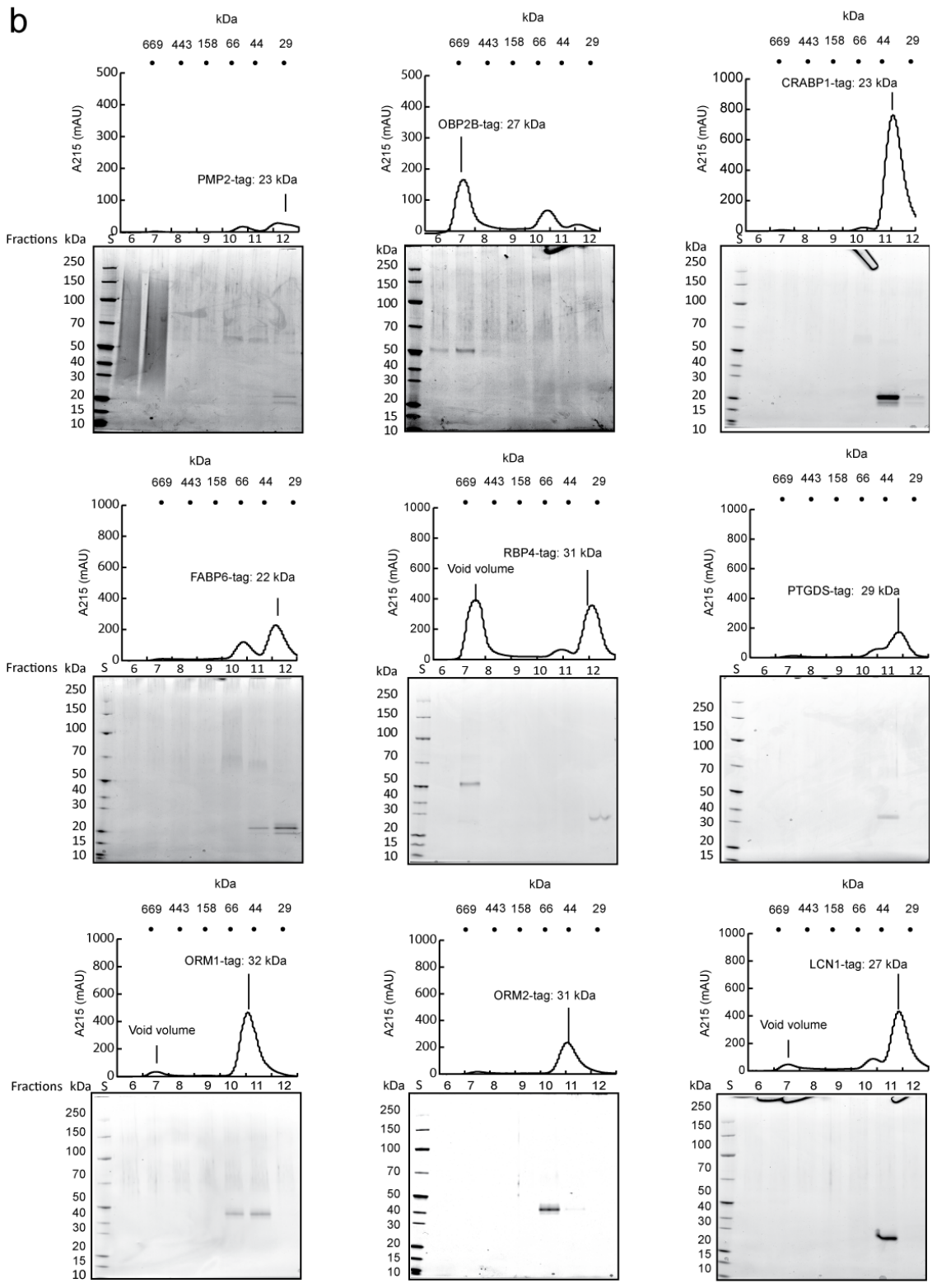

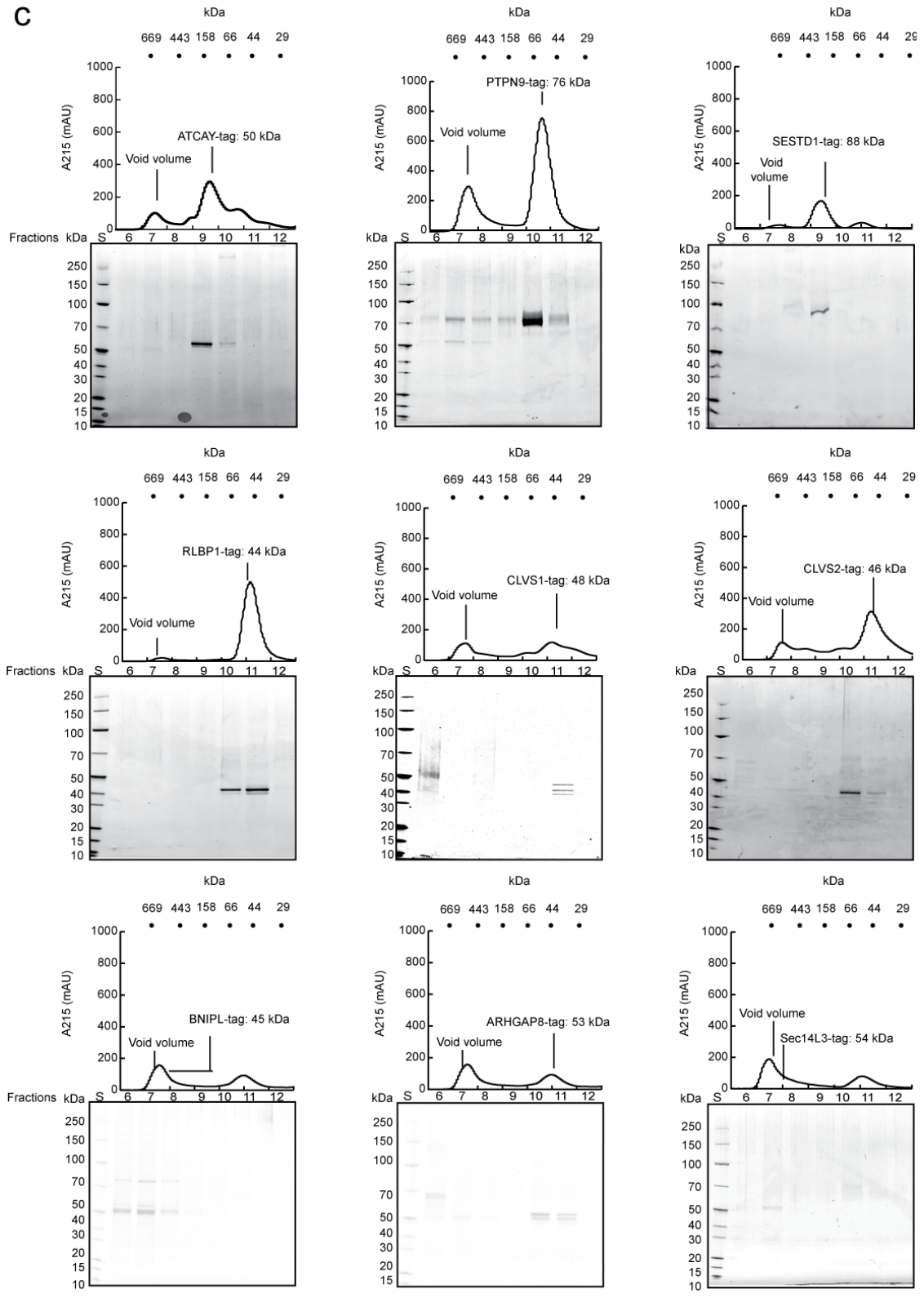

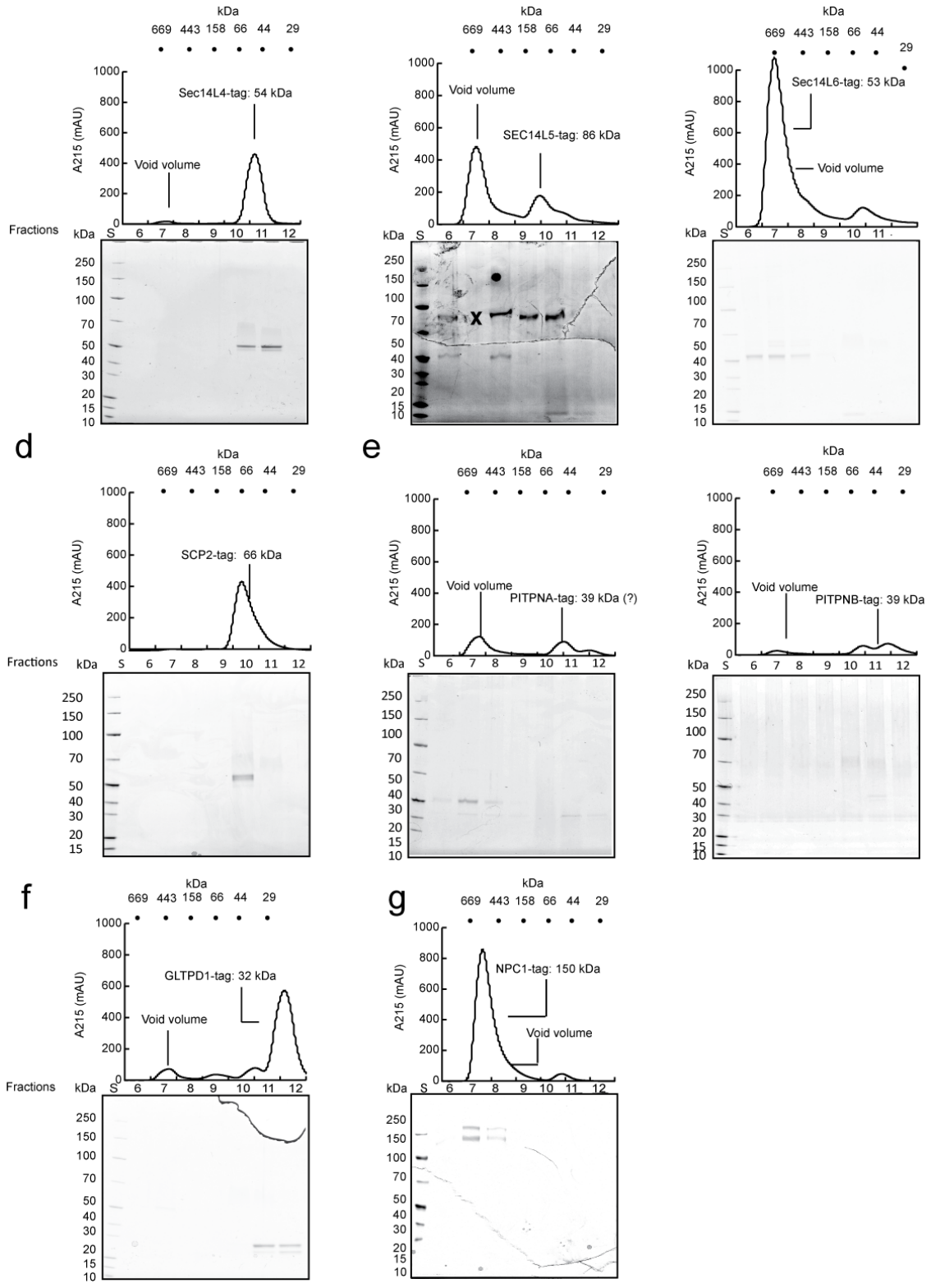

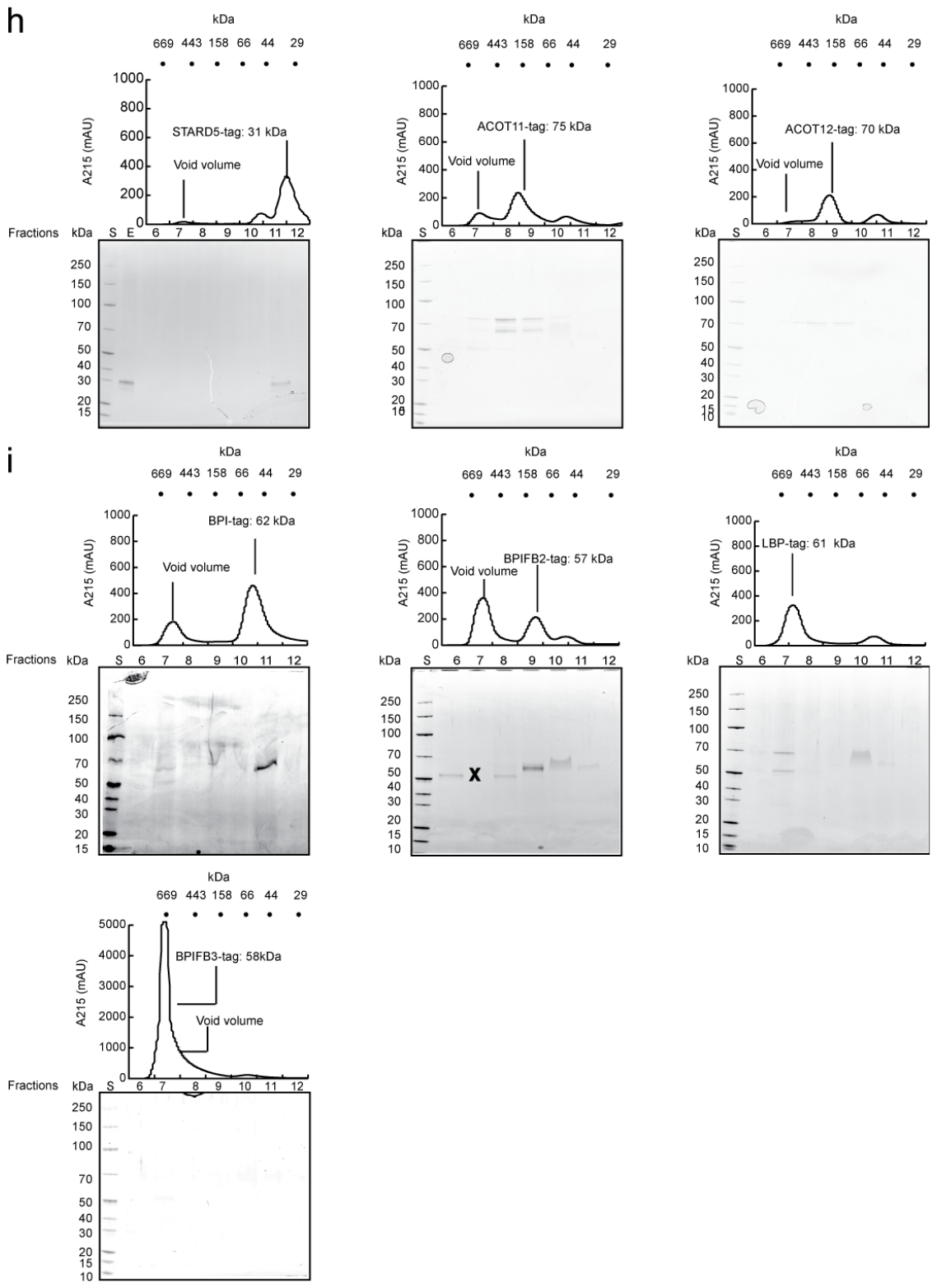

**Supplementary Data S1B:
Purification and characterization of LTPs-lipid complexes *in vitro* (SEC results):** Size exclusion chromatography profiles of *in vitro* purified LTP-lipid complexes. After elution from the affinity column, the LTP-lipid complexes were isolated on a Superdex 200 size exclusion resin packed in a Tricorn 5/150 Column (GE Healthcare).

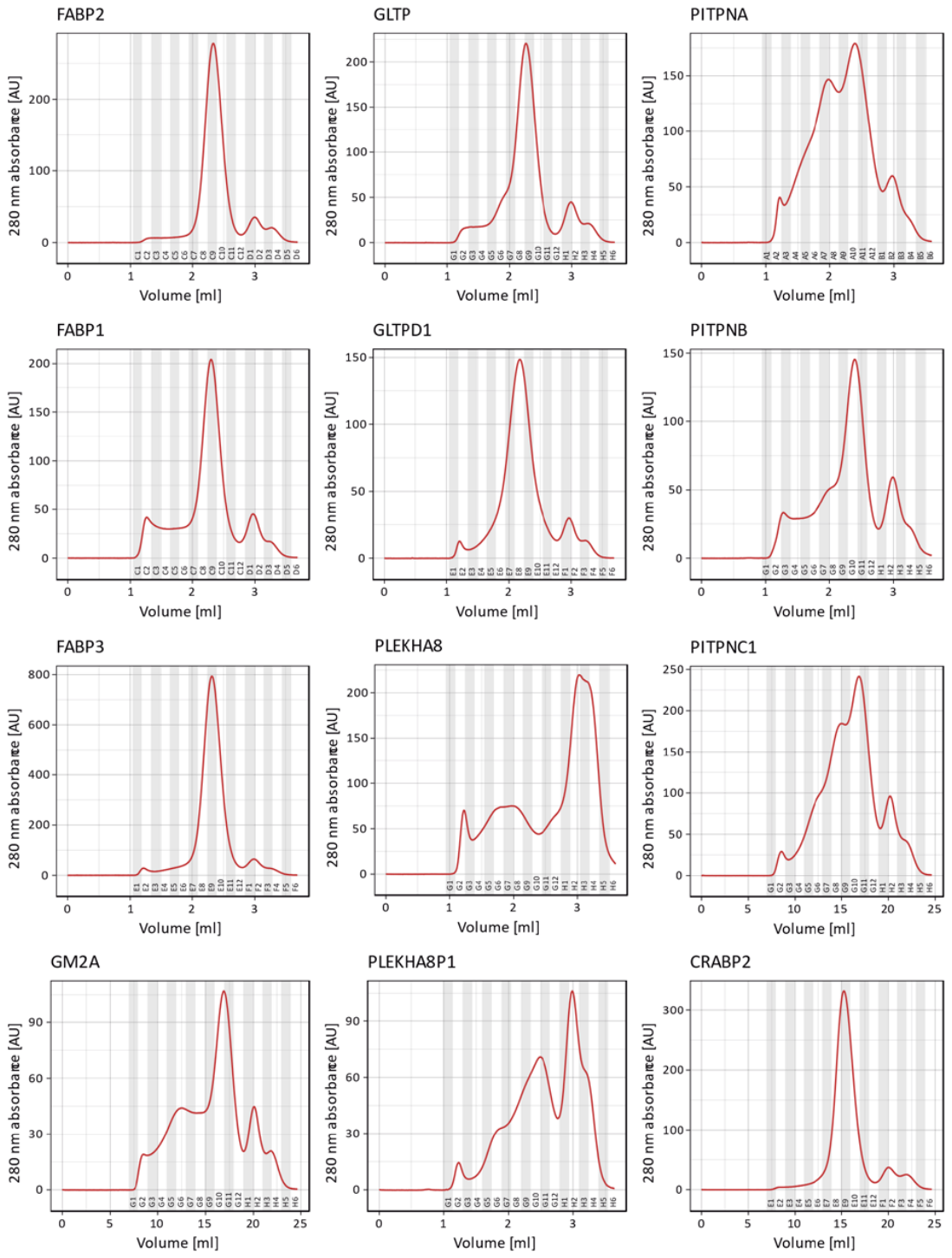

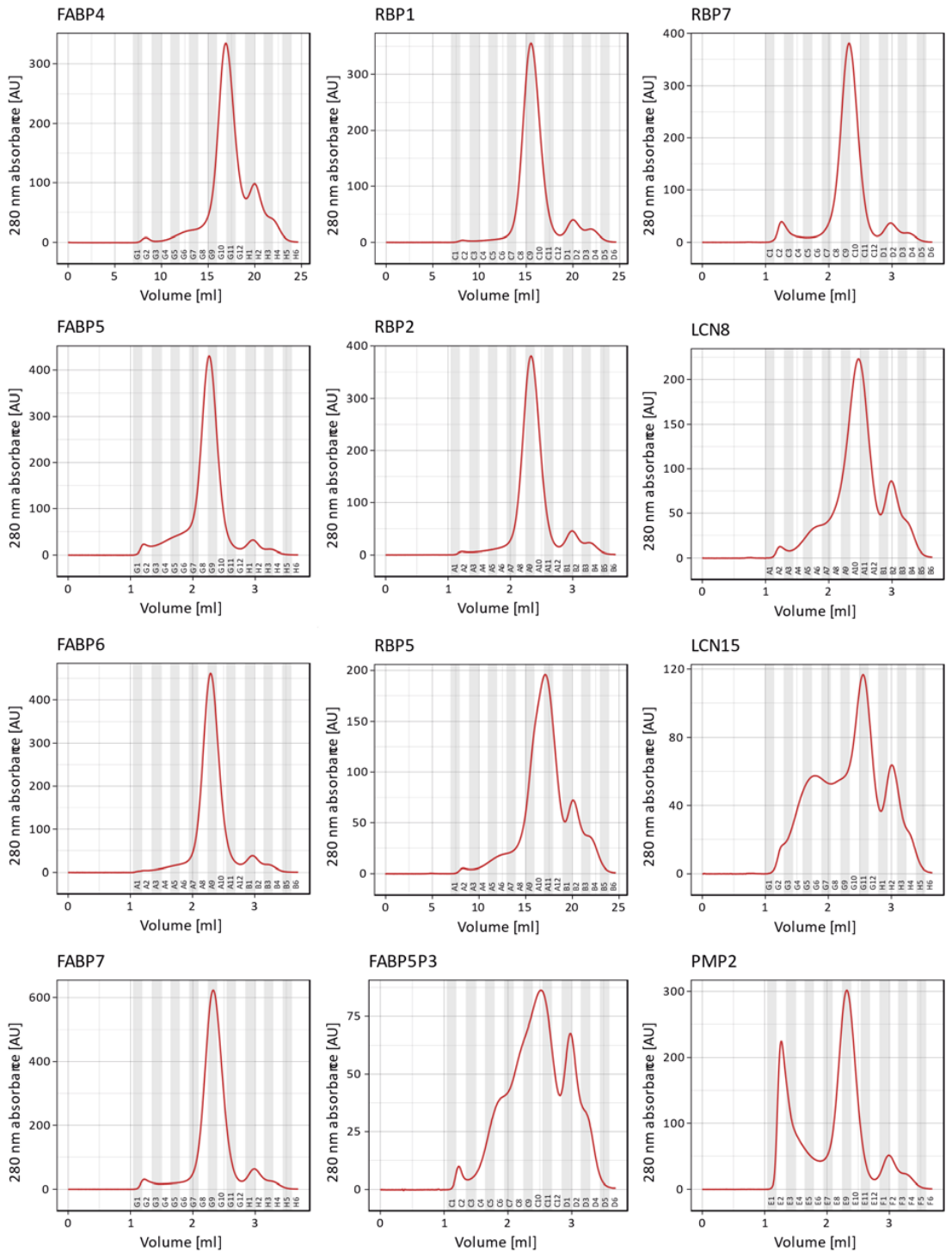

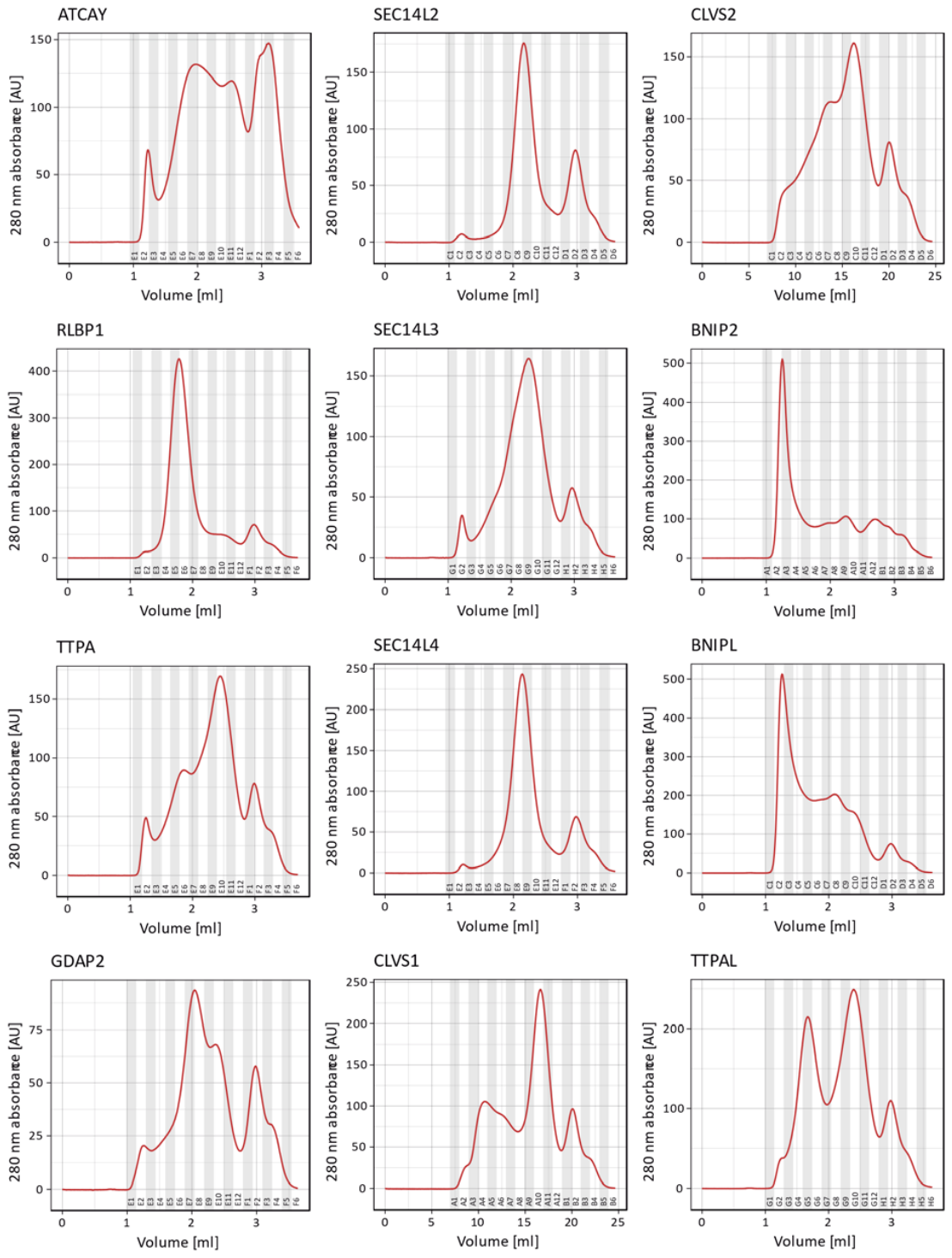

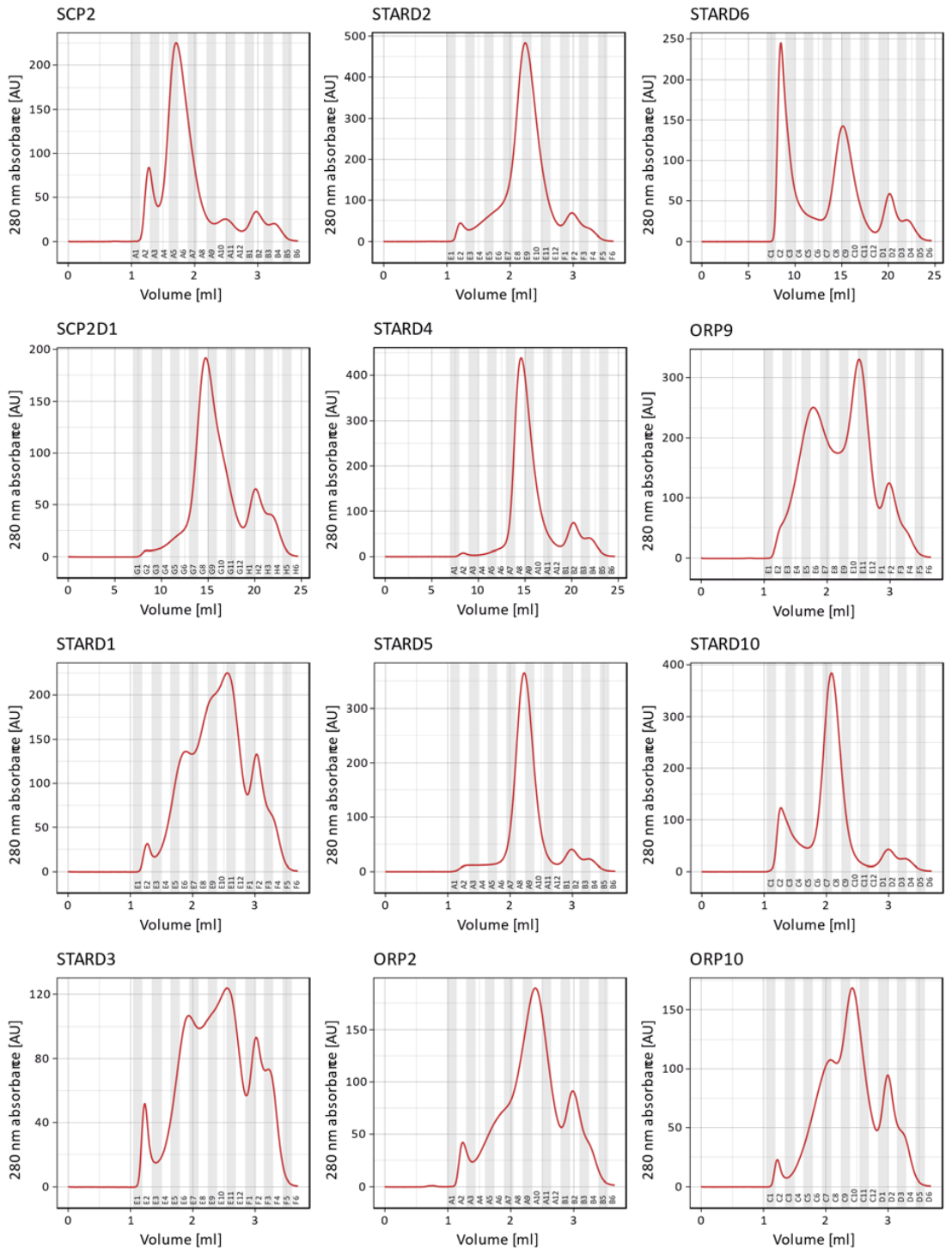

**Purification and characterization of LTPs-lipid complexes *in vitro* (SDS-PAGE results):** SDS-PAGE analysis of SEC fractions of *in vitro* purified LTP-lipid complexes. After elution from the affinity column, the LTP-lipid complexes were isolated on a Superdex 200 size exclusion resin packed in a Tricorn 5/150 Column (GE Healthcare). Proteins gels are stained in colloidal Coomassie Blue.

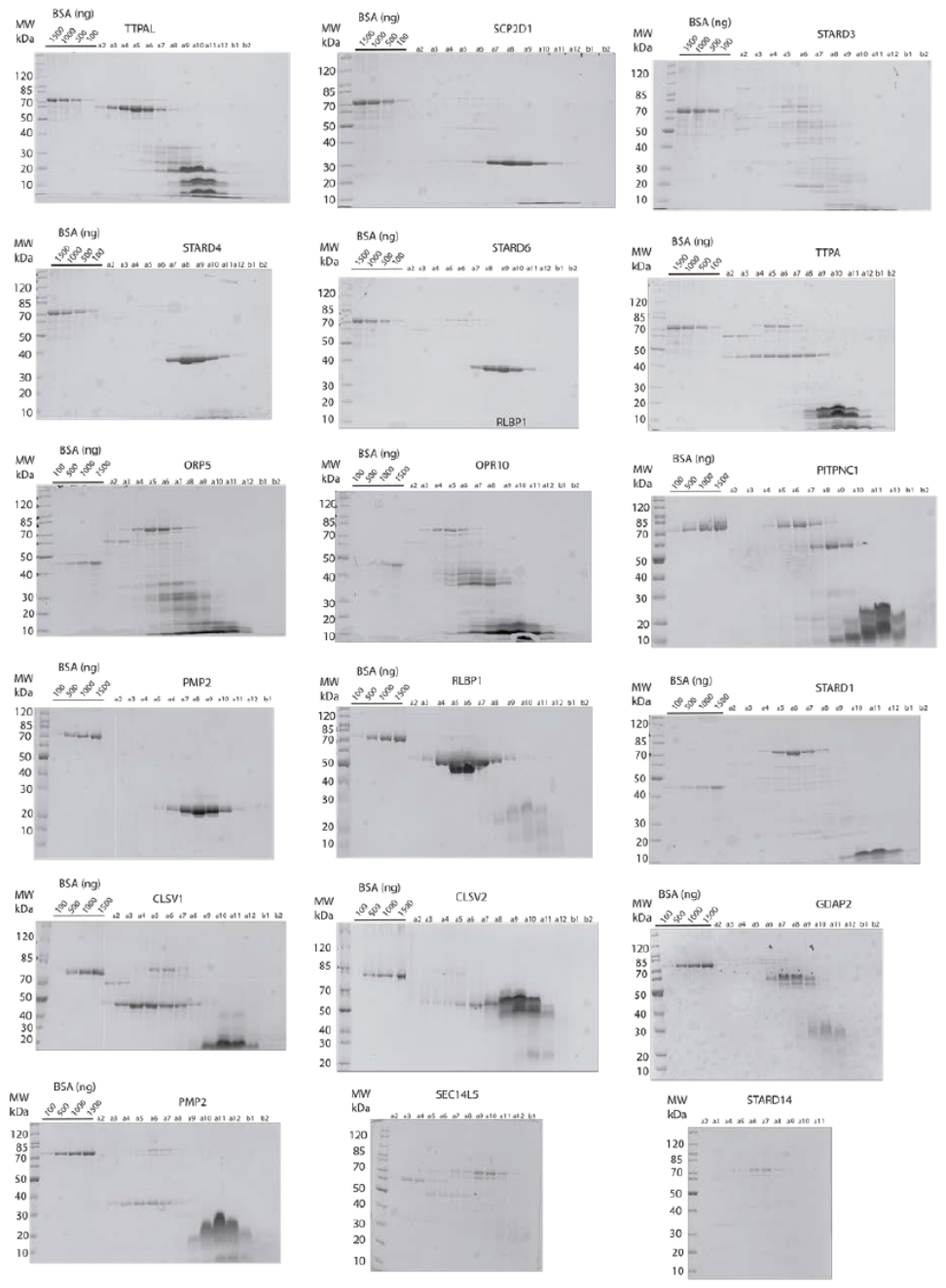
